# HIV-1 Vpu complements HLA-A downregulation by Nef and impairs TCR-mediated recognition of HIV-infected cells through distinct mechanisms

**DOI:** 10.64898/2026.11.04.764327

**Authors:** Mariana G. Bego, Tram N.Q. Pham, Claudia Champagne, Vandana Sharma, Mélanie Laporte, Frédéric Dallaire, Wes Martin, Mark A. Brockman, Frank Kirchhoff, Éric A. Cohen

**Affiliations:** Institut de recherches cliniques de Montréal (IRCM), Montréal, Québec, H2W 1R7, Canada; Department of Microbiology, Infectiology and Immunology, Université de Montréal, Montréal, Québec, H3T 1J4, Canada; Faculty of Health Sciences, Simon Fraser University, Burnaby, British Columbia, V5A 1S6, Canada; Institute of Molecular Virology, Ulm University Medical Center, 89081 Ulm, Germany

**Keywords:** HLA-A2, HIV-1 Vpu primary variants, HLA-A2-Vpu interaction, HLA-A downmodulation, PNC, BST2 restriction, HIV-1 epitope presentation, CD8^+^ T cell anti-HIV response, immune evasion, HIV-1 persistence

## Abstract

Many viruses, including HIV-1, target the human leukocyte antigen (HLA) class I molecules HLA-A, HLA-B, and HLA-C to evade immune surveillance. HIV-1 Nef selectively downregulates HLA-A and HLA-B, whereas Vpu variants from primary strains reduce cell-surface HLA-C expression. Whether HIV-1 also uses Vpu to target other HLA molecules has remained unclear. Here, we show that Vpu from primary strains reduces HLA-A, including the globally prevalent HLA-A2, on infected cells. Although Nef is the principal modulator of these molecules, Vpu acts independently of Nef and impairs TCR-mediated recognition of infected cells in co-culture systems. However, the reduction in TCR signaling is also dependent on the presence of BST2/tetherin, a restriction factor that retains nascent virions and targets them for degradation. HLA-A2-restricted, HIV-specific CTL responses were lowest when both Nef and Vpu were present. Vpu variants from chronic and transmitted/founder strains, but not from the prototypical recombinant strain NL4.3, were sufficient to reduce cell-surface HLA-A2, although activity varied widely among variants and appears to have arisen independently in different hosts. Coimmunoprecipitation and immunofluorescence analyses showed that Vpu interacts with HLA-A2 and redirects it to a perinuclear compartment which is probably the *trans*-Golgi network through a mechanism governed by the Vpu N-terminus/transmembrane domain. Overall, our findings reveal a previously unrecognized function of Vpu and show how HIV-1 uses Nef and Vpu to limit HLA-A-mediated viral antigen presentation and promote immune evasion.

**IMPORTANCE:** We show that the HIV-1 accessory protein Vpu, like Nef, downmodulates HLA-A on infected cells. Because Vpu also downmodulates HLA-C, it can substantially reduce overall HLA class I abundance and, consequently, viral antigen presentation to cytotoxic T lymphocytes (CTLs). Vpu impairs TCR-mediated recognition of infected cells through two distinct mechanisms: HLA-A2 downregulation and reduced virion retention through antagonism of the restriction factor BST2. CTL responses were most diminished when both Nef and Vpu were present, supporting a role for Vpu in evading T cell-mediated immunity. The ability of Vpu proteins from primary HIV-1 strains across clades and groups but not from the prototypical recombinant strain NL4.3 to reduce HLA-A2 levels suggests that HIV-1 evolved an additional mechanism to evade immune recognition and control.

## INTRODUCTION

In people living with HIV, the kinetics of disease progression to AIDS varies greatly from less than a year to several decades (1, 2). The inter-individual variation observed has been linked to host genetic factors that are associated with resistance to HIV infection, slow rate of disease progression, and the likelihood of viral transmission (1, 3, 4). Among others, the *human leukocyte antigen* (*HLA*) class I loci are the genetic variants often linked to the natural history of HIV infection and disease progression (1, 4).The HLA class I (HLA-I) system consists of the classical molecules (HLA-A, -B, and -C), which present peptides to cytotoxic T lymphocytes, and the non-classical molecules (HLA-E, -F, -G and -H), which have more specialized roles in immune regulation and tolerance (5, 6). HLA molecules are highly polymorphic and play a pivotal role in eliciting both innate and adaptive immune responses (5). During viral infections, HLA-I is central to the development of T cell-mediated immunity by presenting immunodominant HIV epitopes to CD8^+^ cytotoxic T lymphocytes (CTLs) (1). The extreme polymorphism of the HLA-I system allows for the presentation of diverse viral peptides (5). For years, HIV-1 was thought to selectively downregulate the highly expressed HLA-A and HLA-B to evade CTL recognition while leaving HLA-C and HLA-E intact (7, 8) in order to benefit from their inhibitory effects on natural killer (NK) cells (9). However, this model was later challenged when studies using HIV-1 primary strains revealed that Vpu reduced HLA-C surface expression (10, 11) and that Nef downregulated HLA-E surface levels (12). It then became evident that the initial model was based on findings from studies using the widely-used prototypical HIV-1 recombinant strain NL4-3 derived from cell culture adapted strains (13),which lacks these abilities (8, 14–16), thus revealing important functional differences between primary isolates and NL4.3-like strains.

HIV-1 Nef and Vpu are highly multifunctional proteins tailored to modulate various host receptors including membrane-associated restriction factors and immune cell markers. The ability of Nef and Vpu to downregulate such proteins is critical for efficient transmission, maintenance, and pathogenicity of HIV-1 *in vivo* (17, 18). However, there is functional redundancy between HIV-1 Nef and Vpu since both proteins can modulate multiple receptors at the surface of infected cells, including the HIV-1 receptor, CD4 (reviewed in (17, 19)). This notable redundancy is likely the result of coevolution of both genes, despite their highly divergent amino acid sequences and the absence of shared functional motifs or membrane topology (20). The *vpu* gene was likely acquired by a common simian ancestor of *vpu*-containing *Cercopithecus* viruses, which later recombined with other simian immunodeficiency viruses (SIVs), to eventually give rise to SIVs infecting chimpanzees and gorillas, and ultimately HIV-1 (21). In contrast, the *nef* gene is found in the genomes of all primate lentiviruses. In most non-*vpu*-containing viruses, Nef was shown to counteract BST2 (tetherin), a known membrane-associated restriction factor specifically antagonized by HIV-1 Vpu, which entraps nascent viral particles and target them for degradation in the endolysosomal pathway (22–28). Nef and Vpu interact with their targets via direct physical association, with Nef binding to the cytoplasmic tail and Vpu, the transmembrane (TM) domain of the targeted host factors (29–33). Nef typically intercepts host targets within receptor transport routes to and from the plasma membrane, while Vpu preferentially targets newly synthesized or recycling cargos (17, 20, 34, 35). In the context of HLA-I, primary HIV-1 strains are proposed to preferentially target HLA-A and HLA-C through the actions of Nef and Vpu, respectively. However, it remained unclear whether Vpu also targets HLA-A to facilitate optimal evasion of the immune system.

In this study, we show that Vpu from primary HIV-1 strains, but not from the prototypical recombinant strain NL4.3, downregulates HLA-A, especially the worldwide common HLA-A2 (36), and this is in part linked to impaired T cell receptor (TCR)-mediated signaling by effector T cells. BST2 depletion by Vpu also contributes to the impairment. HLA-A2-restricted, virus-specific CTL responses were most reduced in the presence of Nef and Vpu. Vpu is sufficient to downmodulate HLA-A2 although the magnitude of the effect varies widely between primary Vpu from different clades and groups. Mechanistically, Vpu interacts with HLA-A2 via its N-terminus/TM domain (TMD) and limits its expression at the cell surface by redirecting HLA-A2 intracellularly. The observation that the NL4.3 Vpu is defective at modulating HLA-A2 and forming complexes with the HLA molecule and modulating HLA-A2 likely explains why this function was overlooked in earlier studies using the prototypical NL4.3 Vpu. Collectively, our findings suggest that HIV-1 has evolved to exploit both Nef and Vpu to target the HLA-I system by multiple mechanisms, thereby avoiding immune recognition, and elimination of infected cells.

## RESULTS

### HIV-1 Vpu contributes to the downregulation of HLA-A2 on infected CD4^+^ T cells via a mechanism unrelated to transcription

Since the selective downregulation of HLA-A by Nef was mainly examined with virus derived from the prototypical infectious recombinant molecular clone NL4.3 that encodes Vpu and Nef from the cell culture adapted strain LAV (13), we assessed whether Vpu proteins from primary viruses can target these molecules, and to what extent the activity was measurable in the presence of Nef. Thus, CD4^+^ T cells from HLA-A2 donors were infected for up to 72h with HIV-1 NL4.3-GFP, NL4.3-ADA-GFP or the primary transmitted/founder (T/F) WITO virus which express the NL4.3, ADA or WITO Vpu variants, respectively. Wild-type (WT) versions of these viruses along with their derivatives lacking Vpu (dU), Nef (dN) or both Vpu and Nef (dUdN) were included in the analysis (Fig. S1A-C). WT viruses were more infectious than most mutant counterparts (Fig. S1C), consistent with the role of Nef and Vpu in infectivity (reviewed in (37, 38)). However, this did not affect the current assessment examining the effect of Vpu on HLA-A2 expression on infected (GFP^+^ or p24^+^) cells. Consistent with the role of Vpu as the principal modulator of BST2 expression, BST2 levels were reduced on WT and dN virus-infected cells (Fig. S1B). CD4 expression was decreased on infected cells (Fig. S1B) in line with the role of Nef, Vpu and Env as viral modulators of CD4.

Using an HLA-A2–specific monoclonal Ab (clone BB7.2), we found that all WT viruses downmodulated HLA-A2 at the surface of infected cells (Fig. 1A-B). Vpu from ADA and WITO viruses, but not from NL4.3, contributed to the downregulation since cells infected with Vpu-deficient viruses consistently expressed significantly higher levels of surface HLA-A2 compared to their Vpu-expressing counterparts (up to two-fold difference when compare WT and dU or dN and dUdN, Fig. 1A-B). Of note, Nef has a major contribution to this process, robustly modulating HLA-A2 expression on infected cells. A similar analysis for HLA-C showed distinct expression profiles on infected cells. The HLA was downregulated efficiently on cells infected with the ADA or WITO, and the effect was entirely dependent on Vpu (Fig. S1D). However, consistent with previous reports (10, 11), no changes were observed in NL4.3-infected cells.

**FIG 1.**
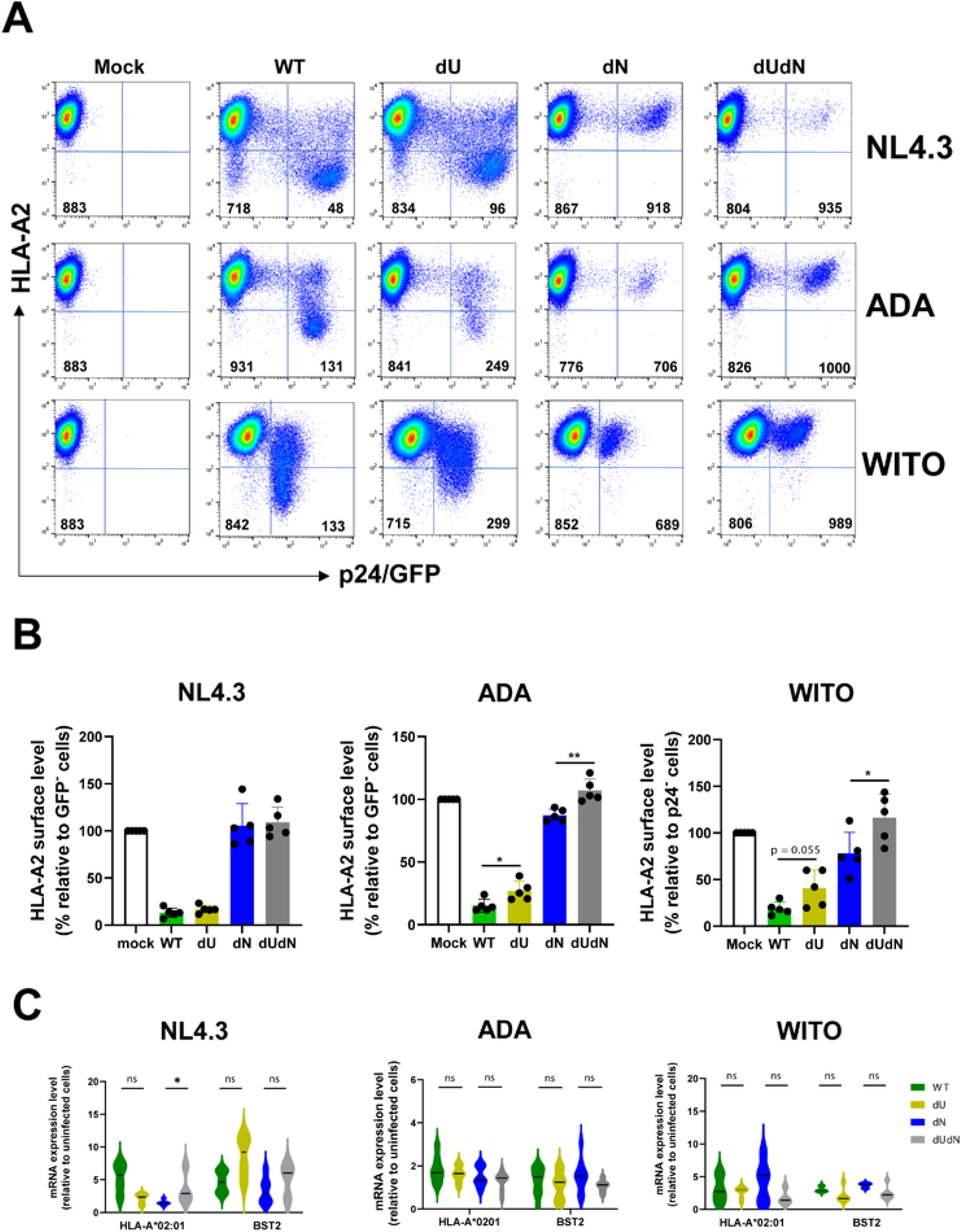
Contribution of HIV-1 Vpu to the modulation of cell-surface HLA-A2 expression on infected CD4^+^ T cells. (**A-B**) Primary CD4^+^ T cells were infected with GFP-marked NL4.3, GFP-marked CCR5-tropic NL4.3-ADA (ADA), or WITO viruses expressing all viral proteins (WT) or lacking Vpu (dU), lacking Nef (dN) or lacking both proteins (dU/dN). Cells not exposed to HIV-1 (mock) were used as negative controls. At 3 days post infection (dpi), samples were analyzed by flow cytometry for HLA-A2 using an α-HLA-A2 Ab. Infected cells were determined based on the expression of GFP (for NL4.3 and ADA viruses) or HIV p24 (for WITO virus). Panel A: HLA-A2 levels at the surface of infected CD4^+^ T cells from an HLA-A*02:01 donor (#232). Geomean fluorescence intensity (gMFI) values for infected (GFP^+^ or HIV-1 p24^+^) or bystander (GFP^-^or HIV-1 p24^-^) cells are as indicated. Panel B: Compiled data from independent experiments with different HLA-A2 donors. Cell-surface HLA-A2 levels were determined as follows. From each infected culture, the gMFI for HLA-A2 on infected cells was divided by that of bystander (uninfected) cells x 100. Shown are mean +/-SEM (n=5, each dot is a donor). Statistical differences between WT and dU or dU and dUdN were determined by Mann-Whitney’s tests. * < 0.05; ** < 0.005. (**C**) Primary CD4^+^ T cells from HLA-A2 positive donors were infected with WITO or GFP-marked NL4.3-based viruses (NL4.3, ADA). At 3 dpi, productively infected cells were sorted and analyzed for mRNA levels of HLA-A*02:01 or BST2 using RT-qPCR. GAPDH was the normalizer. Shown in violin plots is the mRNA expression levels in infected cells relative to uninfected cells (horizontal lines indicate median values; n = 4 experiments with up to 3 donors for each virus; qPCR was done in duplicates/triplicates). Statistical differences between WT and dU or dU and dUdN were determined by Mann-Whitney’s tests; ns, not significant. See also Fig. S1 and S2.

Since Vpu was previously reported to inhibit activation of the transcription factor NF-κB (39, 40), we assessed whether the differential HLA-A2 levels at the surface of infected cells were transcriptionally regulated. Hence, CD4^+^ T cells from HLA-A2 positive donors were infected and productively infected cells (GFP^+^ in the case of NL4.3 and ADA viruses; gp120^+^ in the case of WITO) were subsequently sorted by flow cytometry. The cells were then assessed by qPCR for HLA-A*02:01 mRNA levels. We noted a general increase in mRNA abundance of the *HLA-A* gene in infected cells compared to uninfected cells and the effect of Vpu and Nef on surface HLA-A2 expression could not be attributable to a decrease in gene transcription. This was true for all three viruses (Fig. 1C). As expected, the effect of Vpu on the level of surface BST2 was also not transcriptionally regulated (Fig. 1C). In summary, these results demonstrate that during *bona fide* HIV-1 infection, primary Vpu variants down-modulate HLA-A2 cell surface expression, albeit to a lesser extent than Nef.

### HIV-1 Vpu also targets other HLA-A subtypes on the surface of infected T cells

To assess whether Vpu can also modulate other HLA-A subtypes, we extended the analysis on donors who were identified through genotyping or flow cytometry to be HLA-A positive but HLA-A2 negative (Fig. 2A). lndeed, as seen with HLA-A2, although Nef plays an important role in modulating HLA-A expression, Vpu from both ADA and WITO viruses consistently complement Nef activity to enhance the downregulation. Notably, the effect was observed independently of Nef (Fig. 2B-C). Thus, the overall findings demonstrate that Vpu negatively affects the expression of other HLA-A alleles, not just the -A2.

**FIG. 2.**
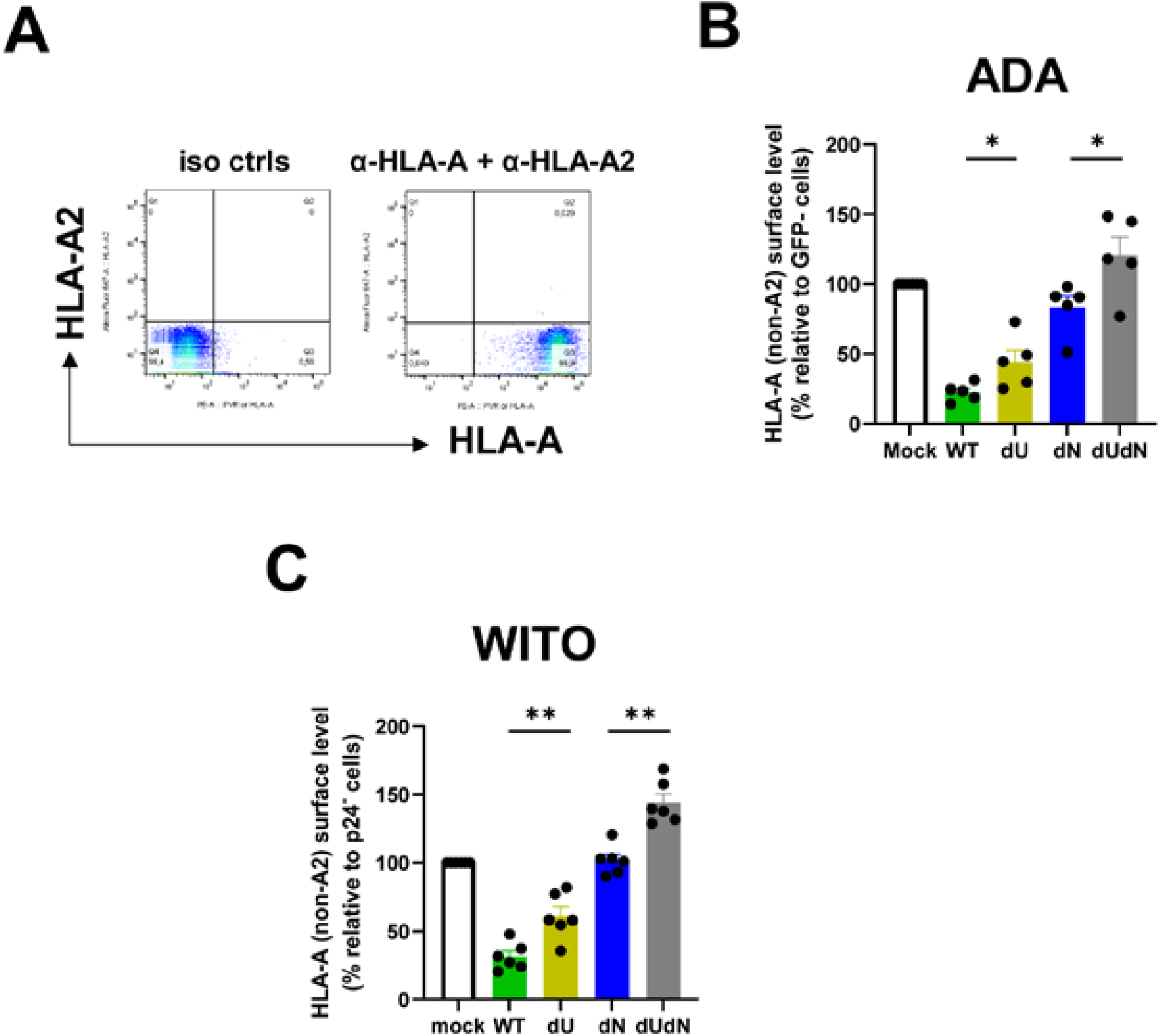
HIV-1 Vpu plays a role in the downregulation of other HLA-A subtypes (non-HLA-A2) at the surface of infected CD4^+^ T cells. (**A)** Identification of HLA-A (non-A2) positive donors by flow cytometry. Uninfected primary CD4^+^T cells were co-stained for HLA-A2 and HLA-A using α-HLA-A2 and α-HLA-A Abs, respectively, or the corresponding isotype controls (iso ctrls). (**B, C**) Primary CD4^+^ T cells from HLA-A (non-A2) positive donors were infected with GFP-marked CCR5-tropic NL4.3-ADA (ADA) or WITO viruses expressing all viral proteins (WT) or lacking Vpu (dU), Nef (dN) or both proteins (dU/dN). Cells not exposed to the virus (mock) were used as controls. At 3 days post infection (dpi), samples were analyzed by flow cytometry for HLA-A using the α-HLA-A Ab as for Panel A. Data analysis and statistical test were as described for Fig. 1. Shown are mean +/- SEM (n=5, each dot is a donor).

### Effect of Vpu-mediated HLA-A2 downregulation on HLA-A2-restricted, virus-specific CD8^+^ T cell responses

To examine the functional significance of Vpu-mediated HLA-A2 downregulation in immune evasion from CD8^+^ T cells, we used a reporter system in which HLA-A*02:01-expressing CEM CD4^+^ T cells (CEM-A2) or parental CEM T cells (Fig. S2A) were infected with NL4.3-ADA-GFP and co-cultured with Jurkat T cells which co-express CD8 and the HLA-A*02:01-restricted, HIV-1 Gag epitope FK10 (FLGKIWPSYK) TCR (Fig. 3A). TCR-mediated effector signaling in response to endogenous FK10 antigen presentation was measured by NFAT-dependent luciferase expression (35). In this system, WT virus reduced HLA-A2 expression on CEM-A2 cells more efficiently than the mutant viruses (17.9% +/-5.1%; Fig. 3B; Fig. S2B), consistent with earlier observations in primary CD4^+^ T cells (Fig. 1). Moreover, the effect of Vpu on HLA-A2 was detectable regardless of whether Nef was there (WT *vs*. dU; dN *vs.* dUdN). Functionally, HLA-A2 levels on infected cells were closely correlated with the magnitude of TCR-mediated effector signaling (Fig. 3C). Importantly, similar results were obtained with WITO, demonstrating that the effect was not strain-dependent (Fig. 3D-E; Fig. S2B). TCR-mediated signaling was not observed when reporter CD8^+^ T cells were exposed to parental HLA-A2 deficient CEM CD4^+^ T cells, supporting the specificity of the signaling (Fig. 3C).

**FIG 3.**
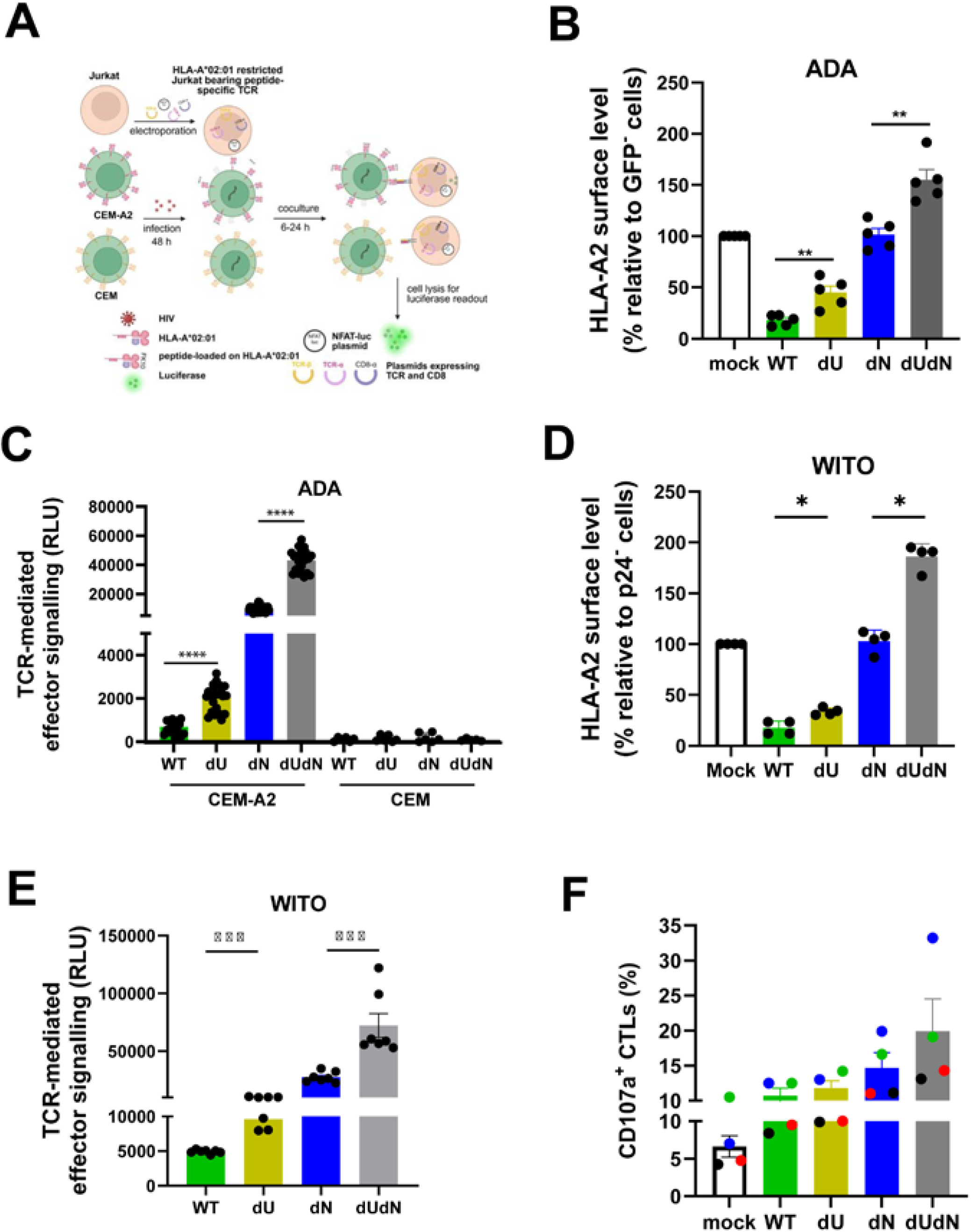
Functional implication of Vpu-mediated HLA-A2 downmodulation on CD8^+^ T cell responses. CEM or CEM-A2 CD4^+^ T cells were infected with wild-type (WT) NL4.3-ADA-GFP or WITO virus along with their respective mutant derivatives deficient of Vpu (dU), Nef (dN) or Vpu and Nef (dUdN) for 48 h. Uninfected cells (mock) were used as a negative control. Thereafter, the CEM T cells were co-cultured with Jurkat reporter cells expressing a TCR specific for the HLA-A*02:01-restricted HIV gag epitope FK10 (FLGKIWPSYK), CD8 and NFAT-driven luciferase for up to 24 h. Samples were analyzed for luciferase expression by bioluminescence. Relative light units (RLU) were a read-out for TCR-mediated signalling of endogenous HIV peptide presentation. A portion of CEM T cells (48 h post infection) was set aside for HLA-A2 analysis by flow cytometry. **(A**) Experimental design. *Created in BioRender. Cai, C.* (*2025*) https://BioRender.com/ciwdl9l. (**B-E)** HLA-A2 at the surface of infected cells were determined as described in Fig. 1 legend. Panels B and D, compiled data over five experiments (mean +/- SEM) for ADA and WITO viruses, respectively. Each dot is an independent infection. Panels C and E, overall results from five different experiments (mean+/- SEM), showing the extent of TCR-mediated signalling following co-cultures of infected cells with CD8-expressing reporter T cells. The assay was done with 2-4 replicates per condition. Each dot represents an individual data point. The *p* values were: * = 0.02; ** = 0.008; *** = 0.0006; **** < 0.0001 (non-parametric Mann-Whitney’s test). (F). CEM-A2 CD4^+^ T cells were infected with WITO viruses expressing all proteins (WT) or deficient for Vpu (dU), Nef (dN) or Vpu and Nef (dUdN) for 24 h. Uninfected (mock) and infected CEM-A2 cells (targets) were subsequently co-cultured for 3h with primary CD8^+^ T cells expressing GFP-marked HIV-based lentiviral vector expressing the HLA-A*02:01 restricted, SL9-specific TCR (effectors). The cytotoxic lymphocytes (CTLs) were analyzed by flow cytometry for CD107a at the cell surface (indicative of cell degranulation). Shown are frequencies (mean ± SD) of Aqua^-^GFP^+^CD8^+^CD107a^+^ CTLs over different donors (each color is a donor). See also Figs. S2 and S3.

To further assess whether HLA-A2 downregulation on infected cells would lead to reduced CTL responses, we quantified the extent of virus-specific, HLA-A*02:01 restricted CD8^+^ T cell responses following co-culture with WITO-infected CEM-A2 cells. Cell surface CD107a, a marker of effector cell degranulation, appeared comparable between WT and dU but was moderately higher with dUdN when compared to dN (Fig. 3F; Fig. S3A). In the same analysis, IFNγ-and TNFα-producing CTLs were most abundant with dUdN and least frequently detected with WT (Fig. S3B-C), although the magnitude of the CTL responses was not closely associated with the extent of virus-induced modulation of HLA-A2 (Fig. 3D) as observed with the reporter CD8+ T cell assay (Fig. 3E).

### BST2 depletion reduces TCR-mediated signaling in CD8^+^ T cells

Vpu antagonizes BST2-mediated virion tethering at the cell surface. We therefore asked whether this activity could indirectly reduce TCR-mediated signaling by potentially decreasing the entrapment and endocytosis of tethered virions and, consequently, antigen presentation. To test this, we repeated the co-culture experiments using BST2-depleted CEM-A2 cells and reporter CD8^+^ T cells (Fig. 4A). BST2 depletion did not affect Vpu-mediated HLA-A2 downregulation (Fig. 4B), but it reduced TCR-mediated signaling relative to the non-targeting (NT) control, particularly in cells infected with WT or Vpu-deficient HIV-1. Moreover, the Vpu-dependent reduction in signaling was abolished when BST2 was knocked down (Fig. 4C; compare WT with dU, or dN with dUdN, under NT and Sh45 conditions). These results suggest that BST2 contributes independently to this process, possibly by promoting the processing and presentation of viral antigens to CD8^+^ T cells. Indeed, it is well established that BST2 restricts HIV-1 release by tethering nascent virions at the plasma membrane and targeting them for degradation in the endolysosomes (26, 28). Consistent with this interpretation, BST2-depleted cells infected with most viruses elicited lower TCR-mediated signaling than BST2-expressing cells infected with the same viruses. Together, these data indicate that both HLA-A2 levels and potentially, antigen-presentation efficiency shape TCR signaling in CD8^+^ T cells.

**FIG. 4.**
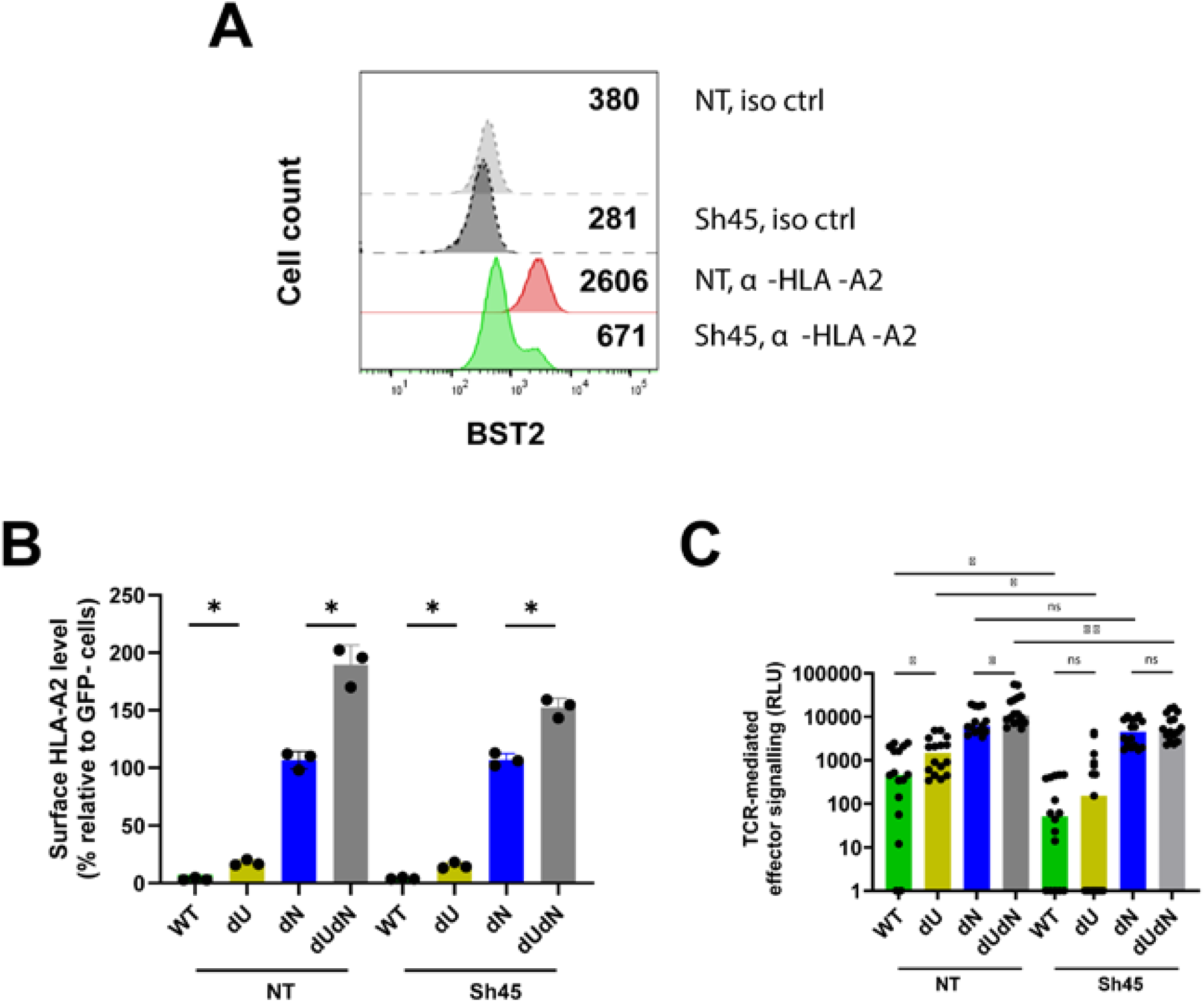
BST2 depletion reduces TCR-mediated signalling in the CD8^+^ T cell reporter system. CEM-A2 transduced with lentiviral vectors expressing nontargeting (NT) or BST2-targetting shRNA (sh45). BST2 depletion was confirmed by flow cytometry using α-BST2 Ab and isotype control (iso ctrl). Panel A, half offset histograms depict BST2 levels with indicated gMFI values. Panels B and C, BST2-expressing or -depleted CEM-A2 were infected with NL4.3-ADA-GFP viruses and examined for HLA-A2 expression as described for Fig. 3B. Infected cells were co-cultured with CD8^+^ Jurkat reporter cells as explained for Fig. 3C. Each dot in Panel B is a different experiment (mean +/- SD). Each dot in Panel C is a data point. Median results from 5 experiments with 2-6 replicates are shown. Statistical test: Mann-Whitney (* < 0.05; ** < 0.005; ns, not significant).

### HIV-1 Vpu alone is sufficient to downregulate cell-surface HLA-A2 expression

We next assessed whether Vpu alone is sufficient to downregulate HLA-A2, and if so, whether the activity is conserved in chronic and T/F HIV-1 strains. For this purpose, we first generated HeLa cells that stably express HA-tagged HLA-A*02:01 (here-after referred to as HeLa-A2 cells) (Fig. S4). Subsequently, HeLa-A2 cells were transfected with an empty GFP-marked plasmid or vectors co-expressing *gfp* and AU1-tagged *vpu* genes and analyzed by flow cytometry for HLA-A2. Vpu from six viruses including three chronic (ADA, 89.6 and JR-CSF) and three T/F (WITO, REJO and CH077) downmodulated HLA-A2 to levels that were statistically different from that of the NL4.3 Vpu (Fig. 5A, Fig. S5A). The other Vpu variants (one chronic and three T/F) induced variable decrease in HLA-A2 in different experiments (Fig. 5A).

**FIG 5.**
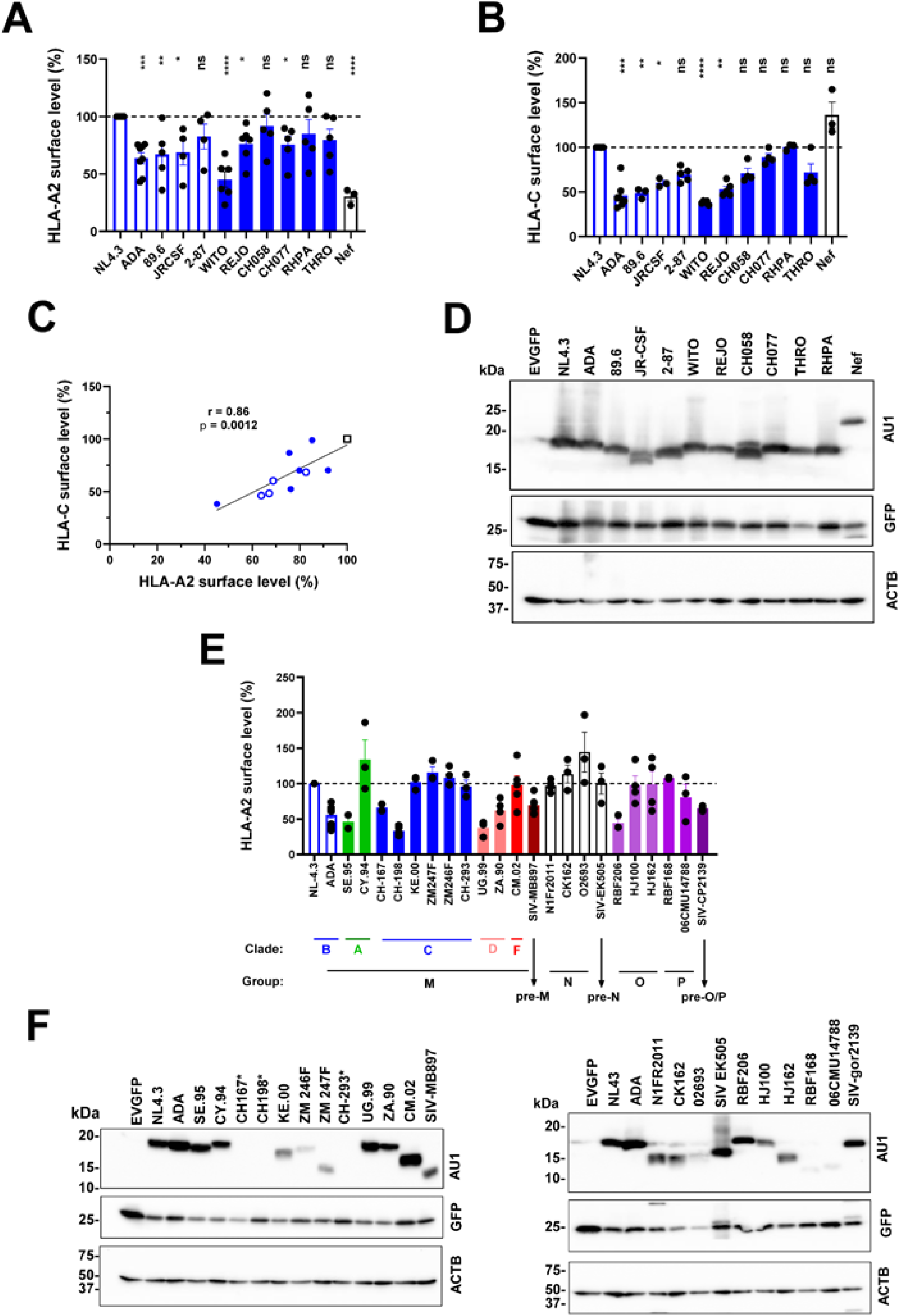
The capacity of Vpu to downmodulate cell-surface HLA-A2 is conserved across distinct HIV-1 clades and groups. HeLa-A2 cells were transfected with a GFP-expressing plasmid encoding either no Vpu or an AU1-tagged Vpu from different primary viruses. At 48 h post-transfection, cell-surface HLA-A2 or HLA-C was analyzed by flow cytometry using α-HLA-A2 or α-HLA-C Abs, respectively. (**A-D**) HeLa-A2 cells expressing Vpu from prototypic NL4.3, chronic, or transmitted/founder viruses of clade B, group M, were analyzed. HIV-1 Nef was derived from NL4.3. Panels A and B show surface HLA-A2 or HLA-C levels on GFP^+^ cells. For each condition, the gMFI of the indicated protein on GFP^+^ cells was divided by that on GFP^-^ cells. Within each experiment, the value obtained with NL4.3 Vpu was set to 100% and used as the reference for all other Vpu variants. Data are shown as mean ± SEM (n = 3 to 5), with each dot representing one experiment. Panel C shows the correlation between the two Vpu activities presented in Panels A and B. Open circles indicate Vpu proteins from chronic viruses, filled circles indicate Vpu proteins from T/F viruses, and the open square indicates NL4.3 Vpu. Correlations were assessed using two-tailed Spearman analyses. Panel D shows Western blot analysis of cell lysates (n = 2). Lysates were probed for AU1 (Vpu), GFP (transfection efficiency), or actin (ACTB; loading control). (**E-F**) HeLa-A2 cells expressing Vpu from different group M clades, selected viruses from groups N, O, and P, and their respective SIV precursors were analyzed. For Panel E, data analysis and graphical presentation were performed as described for Panels A and B. Panel F shows Western blot analysis of cell lysates (n = 2). Untagged Vpu alleles are indicated with an asterisk. In all relevant panels, nonparametric Kruskal-Wallis with Dunn’s multiple comparisons were used, whenever possible, to assess if the differences were statistically significant. The *p* values were: * < 0.05; ** = 0.009; *** = 0.0006; **** < 0.0001; ns, not significant. See also Figs. S4-S7.

Similar analysis for HLA-C (HeLa-A2 cells endogenously express HLA-C*12:03 (41) revealed that Vpu from five of the same six viruses was very efficient, reducing the level by as much as 60% compared to the NL4.3 Vpu (Fig. 5B, Fig. S5B). Indeed, we observed a positive correlation between these two functions (Fig. 5C). Importantly, the lack of HLA-A2 modulation could not be attributable to lack of expression since all variants were well expressed (Fig. 5D). In conclusion, HIV-1 Vpu alone is sufficient to modulate HLA-A2, and the function is conserved in chronic and T/F viruses from clade B of group M, albeit at different levels.

### Vpu proteins from diverse HIV-1 groups, SIVcpz, and SIVgor downmodulate HLA-A2

To assess whether the ability of Vpu to downregulate surface HLA-A2 extends to non-clade B Vpu proteins, we performed the same analysis with a larger panel of Vpu variants. It is evident that Vpu proteins from most clades of group M downregulated HLA-A2, except clade F albeit there was only one Vpu available for testing (Fig. 5E). In addition, 2/6 Vpu C variants (CH-167 and CH-198) showed robust downmodulation (Fig. 5E) and the lack of activity by CH-293 Vpu could not be attributed to the lack of expression since the protein can efficiently downregulate BST2 (Fig. S6).

To gain an insight in the evolution of this function, we extended the analysis to include Vpu variants from SIVcpz*Ptt* strain MB897, the closest genetic relative of HIV-1 group M (42) SIVmus/mon/gsn clade, the original source of Vpu and direct precursor/ancestor of SIVcpzPtt and HIV-1 (43, 44). Here we found that none of the Vpu proteins from the SIVmus/mon/gsn clade modulated HLA-A2 expression but the one from SIVcpz*Ptt* MB897 possessed some activity (mean 70% +/-12% *vs*. 100% for NL4.3 Vpu), suggesting that this function was not present during the evolution of the SIVs in their natural hosts, *mustached* monkeys, *mona* monkeys and *greater spot-nosed* monkeys (Fig. S7A-B). Rather, it was likely acquired prior to cross-species transmission of SIVcpz*Ptt* MB897 and emergence of the pandemic HIV-1.

None of the tested Vpu proteins from group N or from SIVcpz*Ptt* EK505, the closest genetic relative of this group (42) downregulated HLA-A2, suggesting that the function had been acquired independently in different hosts. For the three Vpu O proteins, robust downmodulation was only observed with RBF206, while we detected some activity in 06CMU14788 among the two Vpu P variants tested (Fig. 5F). Interestingly, Vpu from SIVgor CP2139 (45), the closest genetic relative of groups O and P, displays some capacity to downregulate HLA-A2 (Fig. 5E). In summary, the ability of Vpu to reduce HLA-A2 expression extends to non-clade B viruses. However, the function is not conserved in all primary strains tested in that although the activity is observable in certain viruses from groups O and P and the respective SIV precursors, it is not detectable in either group N or its precursor.

Since HeLa cells also express endogenous HLA-A alleles (Fig. S4C), we asked whether Vpu preferentially downregulates these over the stably expressed HLA-A2. Thus, Vpu variants that were deficient/less active were evaluated for their ability to modulate the endogenous HLA-A. It was clear that the ADA and RBF206 Vpu proteins which downregulate HLA-A2 on HeLa-A2 also efficiently decrease HLA-A surface expression in HeLa cells, whereas those unable to alter HLA-A2 expression are also defective at modulating HLA-A (Fig. S7C). Thus, the results strongly exclude the possibility of preferential modulation of endogenous HLA-A alleles by Vpu as the reason for the lack of HLA-A2 downmodulation activity of specific Vpu variants, at least in this cell model. The findings are also in line with the results in primary CD4^+^ T cells which show that Vpu contributes to HLA-A downregulation on infected cells (Fig. 2).

### The N-terminus/TMD of Vpu is critical for downmodulation of HLA-A2

Given the positive correlation between the effect of Vpu on HLA-A2 and HLA-C (Fig. 5C) and the critical role of the N-terminus/TMD of Vpu (Fig. 6A) in HLA-C downregulation (10, 11), we asked if this domain (Fig. S8) could also be important for HLA-A2 downmodulation. Therefore, we generated chimeras for ADA and NL4.3 Vpu such that the ADA-NL Vpu construct would express the N-terminus/TMD of ADA and the C-terminal region of NL4.3, whereas the NL-ADA Vpu was the reverse (Fig. 6B). Robust expression of the Vpu chimeras was verified by Western blot (Fig. 6C). While the ADA-NL Vpu chimera reduced HLA-A2 expression as efficiently as the ADA WT (ADA), the NL-ADA chimera was inactive, indicating that the N-term/TMD confers ADA Vpu the ability to downregulate cell-surface HLA-A2 (Fig. 6D). Analysis for HLA-C showed comparable results (Fig. 6E).

**FIG 6.**
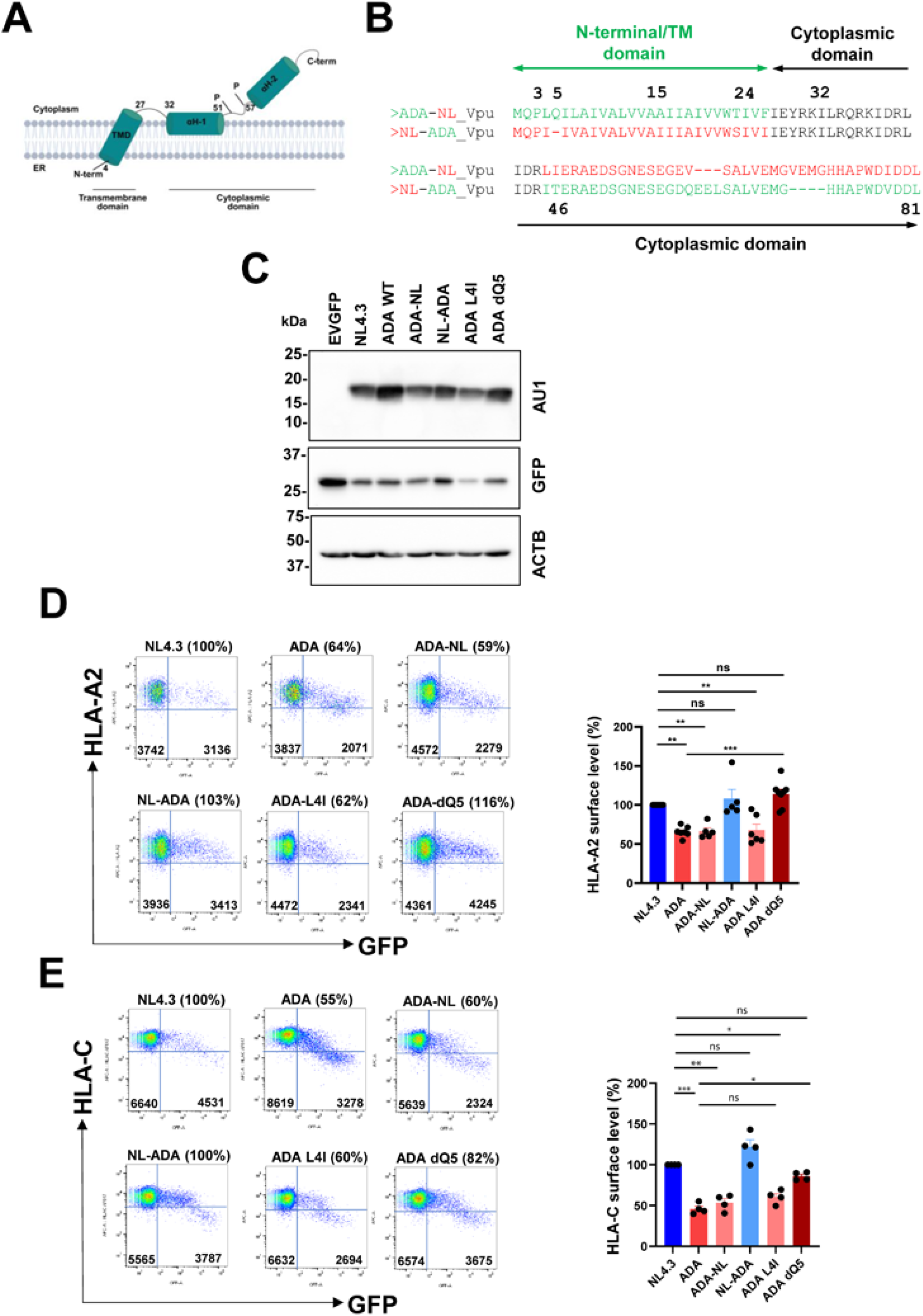
Critical involvement of the N-terminus/TMD of ADA Vpu in the downregulation of HLA-A2. **(A)** HIV-1 Vpu topology depicting the different domains, structural features (alpha helix (αH)) and phosphoserine (P) positions (52 and 56) important for various Vpu functions. *Created in BioRender. Cai, C.* (*2025*) https://BioRender.com/ee7y2mz. (**B)** Sequence of Vpu chimeras generated around the central conserved region (amino acid letters in black) of ADA and NL4.3 Vpu. The ADA-NL chimera contains the N-terminus/TMD of ADA and the C-terminal domain of NL4.3, while the NL-ADA chimera is the reverse. ADA sequence is in green, and NL4.3 in red. Numbering was based on the ADA Vpu sequence. (**C-E**) HeLa-A2 cells were transfected for up to 48 h with a GFP-harbouring plasmid expressing AU1-tagged Vpu. Cells were analyzed for surface HLA-A2 or HLA-C using flow cytometry as described in previous Figures or protein expression by Western blotting. Panel C: Western blot analysis of cell lysates (n=2). EVGFP, empty vector. Panels D and E: cell-surface level of HLA-A2 or HLA-C on GFP^+^ cells. The Abs used to detect HLA-A2 and HLA-C were as described for Fig. 4. For each condition, the gMFI for HLA-A2 or HLA-C on GFP^+^ cells was divided by that on GFP^-^ cells. Then, within each experiment, the value obtained with the NL4.3 Vpu was set as 100% (reference), and all other conditions were compared to this reference. Shown are mean +/- SEM (n = 4 to 8). Fow cytometry dot plots were from one of the experiments. Nonparametric Kruskal-Wallis with Dunn’s multiple comparisons were used to evaluate statistical significance; * < 0.05; ** =0.009; *** < 0.0005; ns, not significant. See also Fig. S8-S9.

Considering the stark contrast in the type of amino acids at positions 4 and 5 between active ADA Vpu and inactive NL4.3 Vpu (Fig. 6B), we asked whether changes at these positions would alter the ability of Vpu to modulate HLA-A2. To this end, we found that the L4I substitution alone did not affect the ability of ADA Vpu to modulate HLA-A2 expression, but deleting the glutamine at position 5 completely abolished the function (Fig. 6C-D). Similarly, the ADA L4I Vpu still downregulated HLA-C while the ADA dQ5 Vpu lost this function (Fig. 6E). Previous data showed that substituting the leucine with an isoleucine at position 4 and combining it with a deletion of the glutamic acid at position 5 impaired the ability of WITO Vpu to downmodulate HLA-C (11).

To further assess the contribution of other N-terminal/TMD residues (Fig. 6B), we swapped the amino acids at position 2 or 6 between the active ADA Vpu and the modestly active 2-87 Vpu (Fig. 5A). Substitutions at positions 2 (Q to K in ADA and K to Q in 2-87) had no impact on the extent of HLA-A2 downmodulation (Fig. S9A and S9B). By contrast, substitution at position 6 in active ADA (I to T) led to a significant reduction in activity, while the inverse mutation in the modestly active 2-87 (T to I) was associated with a meaningful increase in activity (Fig. S9A and S9B). In addition, replacing the threonine in the active REJO Vpu for a proline did not negatively affect the function (Fig. S9C). Together, these results underscore the importance of the identity of neighboring amino acids within the TMD. Natural variations in the sequence of other primary Vpu variants further suggest that certain amino acids at additional positions within this region may also influence the ability of Vpu to downregulate HLA-A2 (Fig. S8).

### Vpu-mediated downregulation of surface HLA-A2 expression is not associated with enhanced internalization

We next assessed by flow cytometry whether the pronounced decrease in HLA-A2 level at the surface is a consequence of accelerated internalization. Thus, HeLa-A2 cells transfected with a Vpu-encoding (or empty) plasmid co-expressing GFP were labelled on ice with an unconjugated mouse α-human HLA-A2 Ab (clone BB7.2) and then chased at different time intervals for up to 4 h at 37°C. The original pool of labelled HLA-A2 on the transfected (GFP^+^) cells was revealed using a fluorescently labelled α-mouse secondary Ab (Fig. 7A). In this experiment, we found no difference in the rate of HLA-A2 internalization between Vpu-positive and Vpu-negative cells (Fig. 7B), suggesting that the heightened decrease in HLA-A2 level when ADA Vpu was present is not a consequence of enhanced internalization.

**FIG 7.**
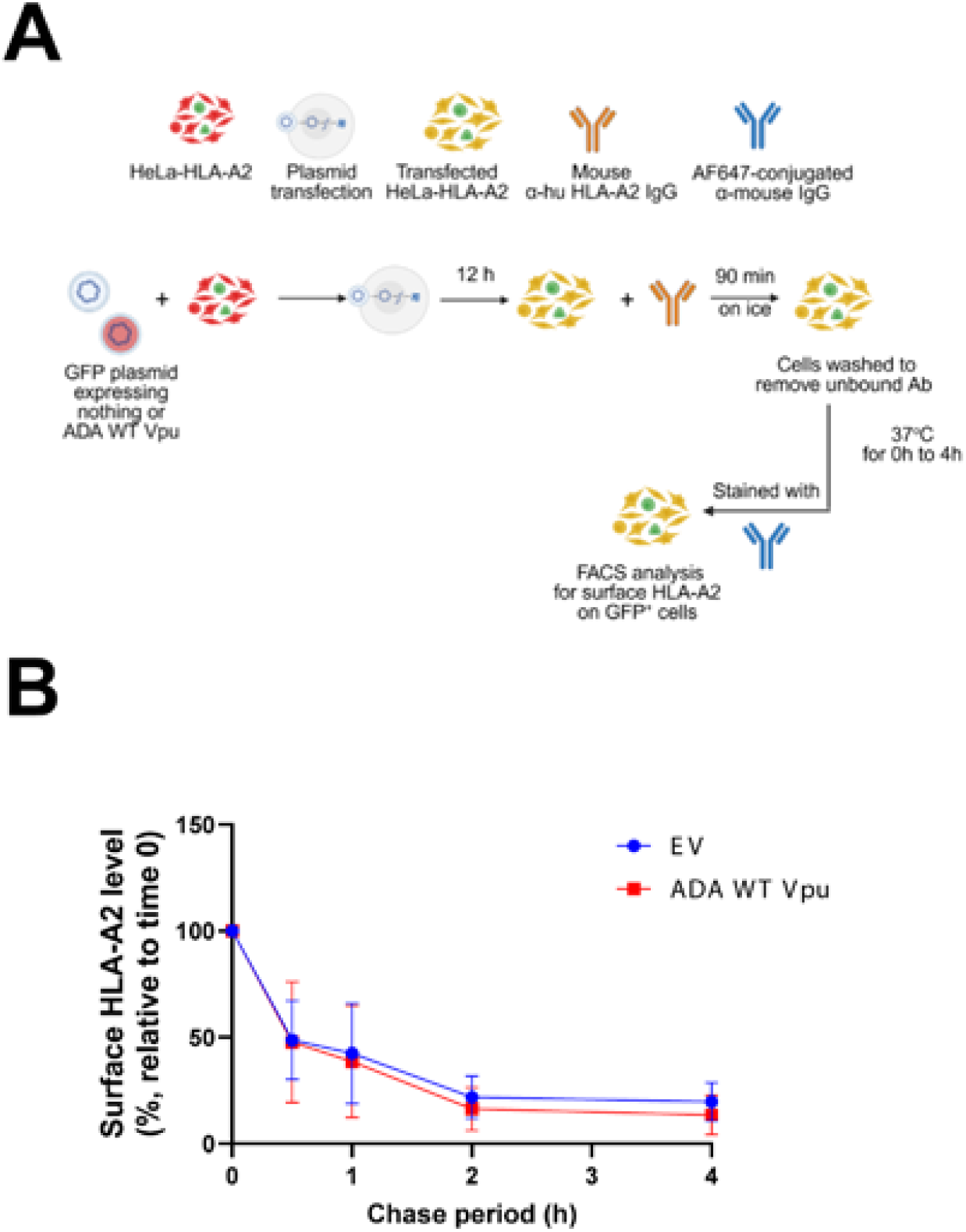
ADA Vpu-mediated downregulation of surface HLA-A2 is not associated with enhanced internalization of HLA-A2. **(A-B)** HeLa-A2 cells were transfected with a GFP-harbouring plasmid expressing no Vpu (EV) or AU-tagged ADA Vpu WT. At 12 h post transfection, cells were labelled with unconjugated mouse α-human HLA-A2 Ab on ice for 90 min and subjected to a 4-hour chase at 37°C. At indicated time points, cells were surface stained with AF647-conjugated α-mouse Ab and analyzed by flow cytometry for the disappearance of the labelled HLA-A2 from the cell surface. Panel A depicts the experimental design. *Created in BioRender. Cai, C.* (*2025*) https://BioRender.com/4ti997n. Panel B shows surface HLA-A2 level (mean +/- SD, n=3) over the 4-hour chase at 37°C relative to time 0 (start of the chase). For each condition (EV or ADA WT), the gMFI for HLA-A2 on GFP^+^ cells at time 0 was set at 100% (reference). HLA-A2 levels at subsequent time points were expressed as percentages of the respective reference (mean +/- SD, n = 3).

### HLA-A2 co-localizes with Vpu in a perinuclear compartment

To further understand the molecular mechanism by which ADA Vpu downmodulates HLA-A2 level at the cell surface, we evaluated the subcellular localization of HLA-A2 relative to Vpu and determined whether these proteins co-localize within a specific compartment using confocal microscopy. In this analysis, HeLa-A2 cells expressing just the EV (no Vpu) or different versions of Vpu were co-stained with α-AU1 Ab (to reveal Vpu) or α-HA (to reveal HLA-A2). The cells were counterstained with Hoechst 33342 to reveal the nuclei. As shown in Fig. 8A, HLA-A2 (red staining) was found mostly at the surface of Vpu-negative cells (blue arrows in Panels i to iv). In ADA WT Vpu-positive cells (white arrows in Panels i to iii), surface HLA-A2 was reduced compared to Vpu-negative cells in the same field (Panel ii, blue *vs*. white arrows). However, no such differences were readily observed in cells expressing the NL4.3 or ADA dQ5 Vpu (Panels i and iii; blue and white arrow comparisons within each Panel). These results were consistent with flow-cytometry data showing that ADA WT Vpu-expressing cells had lower levels of cell-surface HLA-A2 compared to cells expressing ADA dQ5 Vpu or NL4.3 Vpu (Fig. 6).

**FIG 8.**
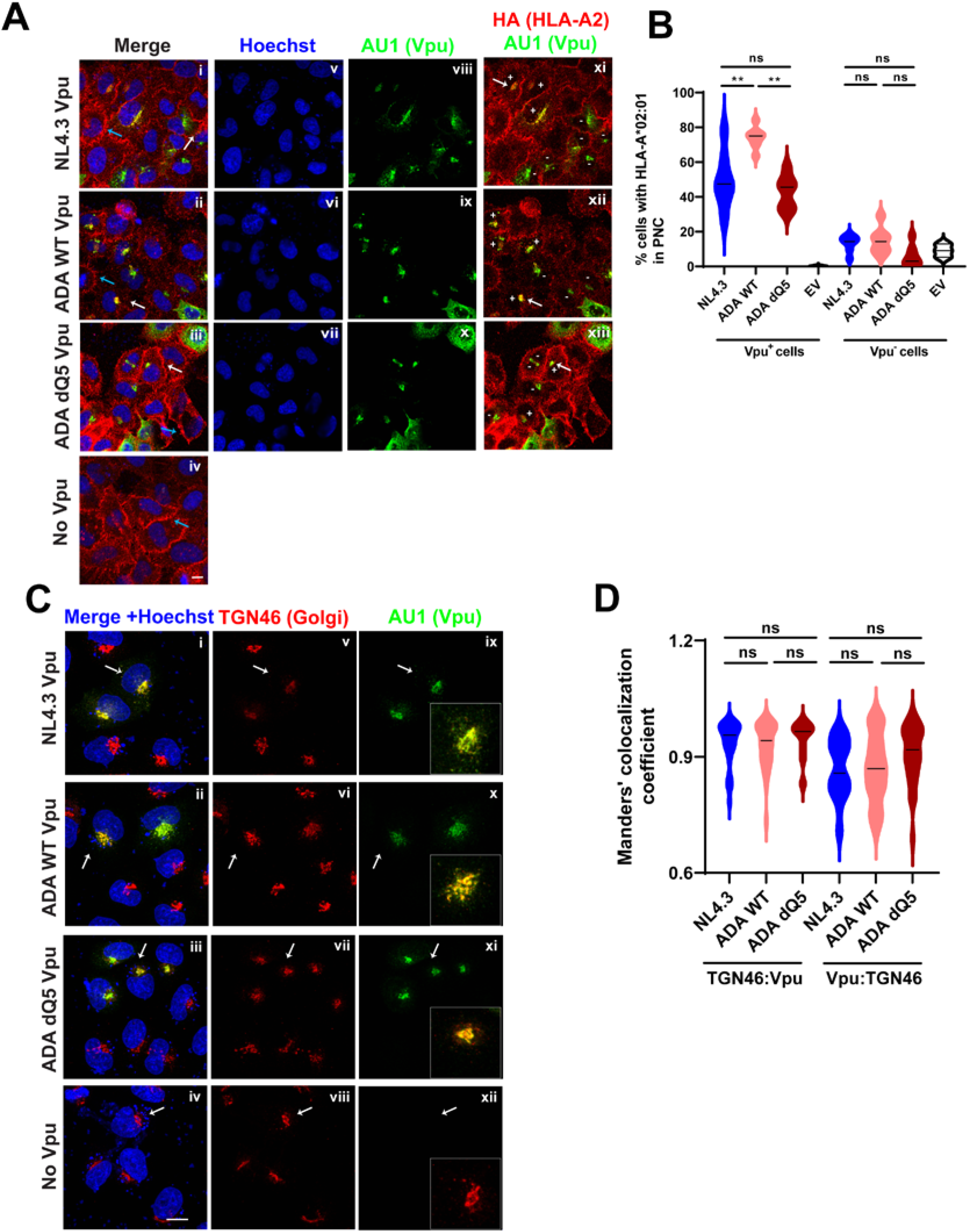
Vpu induces a redistribution of HLA-A2 in a Vpu-containing perinuclear compartment that co-localizes with the *trans*-Golgi network marker TGN46. HeLa- A2 cells stably expressing HA-tagged HLA-A*02:01 were transfected with an empty plasmid (no Vpu), or a construct that encodes the AU1-tagged NL4.3, ADA WT or ADA dQ5 Vpu. Cells were co-stained with α-AU1 (green) and α-HA.11 (red) or α-TGN46 (red) Abs to detect Vpu, HLA-A2 and TGN, respectively. Nuclei were counterstained with Hoechst 33342 (blue). Cells were observed by confocal microscopy, and images captured with a 63x objective lens. (**A)** Representative single-plane images of cells. Panels i to iv: superimposed images of the red, green and blue staining. Blue and white arrows point to HLA-A2 staining in Vpu-negative cells and Vpu-positive cells, respectively. Panels v to vii: Hoechst staining reveals cell nuclei. Panel viii to x: Vpu staining as determined by AU1 signal. Panel xi to xiii: Vpu-positive cells with HLA-A2 co-distributed in a perinuclear compartment (PNC) are marked with the + sign, while those without HLA-A2 in the PNC are denoted by the - sign. The white arrows point to the location of HLA-A2 in a Vpu-containing PNC. (**B**) Percentages of cells with HLA-A2 in PNC (mean +/- SEM) following quantification of multiple images like that shown in Panel A. For each condition, over 100 AU^+^ cells and 200 AU^-^ cells were counted. (**C)** Representative image stacks of Vpu co-localized with the TGN marker TGN46. In all panels, white arrows denote the cell with highly noticeable TGN46-Vpu colocalization. Panels i to iv: superimposed images of the red, green and blue staining. Panels v to viii: TGN46 staining of the Golgi. Panel ix to xii: Vpu staining as determined by AU1 expression. The white squares show an enlarged version of the cell depicted with the white arrow for better viewing of the TGN46 and Vpu co-distribution. (**D**) Co-localization of TGN46 and Vpu as determined by Manders’ coefficient analysis using the ‘colocalization threshold’ plugin in ImageJ. The analysis was performed to assess the probability of TGN46 overlapping with Vpu (TGN46:Vpu), and *vice versa* (Vpu:TGN46). Over 40 cells were observed over multiple fields of view for each Vpu variant. In Panels A and C, scale bar is 10 µm. In Panels B and D, statistical tests used were non-parametric Kruskal-Wallis with Dunn’s multiple comparisons. *P* value: ** < 0.005; ns, not significant.

Based on earlier findings (Fig. 5, Fig. 6), we investigated whether HLA-A2 subcellular distribution differed among cells expressing the different Vpu variants. As illustrated in Fig. 8A, HLA-A2 was found in a perinuclear-like compartment (PNC) when Vpu was also present, and this was observed regardless of the Vpu (Panels xi to xiii, white arrows point to the area of co-distribution, and cell marked with the ‘+’ sign depict HLA-A2 and Vpu co-distribution). Furthermore, HLA-A2 was found more frequently in the PNC in ADA WT Vpu-expressing cells (4/7 in Panel xii, Fig. 8A and Fig. 8B), compared to that in NL4.3-positive cells (3/9 in Panel xi, Fig. 8A and Fig. 8B) or ADA dQ5 Vpu-expressing cells (2/6 in Panel xiii, Fig. 8A and Fig. 8B). Similar comparisons of HLA-A2 subcellular localization were made for Vpu-negative cells, and the level of HLA-A2 in the PNC was markedly lower (Fig. 8B, Vpu-cells).

Analysis for subcellular localization of Vpu showed that all three Vpu variants colocalized with TGN46, as shown by the overlap of the red (TGN46, Fig. 8C) and green (AU1-Vpu, Fig. 8C) signals in the merge panels (Fig. 8C, Panels i-iii). These data are consistent with previous reports demonstrating that Vpu is localized in the perinuclear compartment corresponding to the TGN (33, 46). As well, the degree of colocalization between Vpu and the TGN marker was comparable between cells expressing the ADA WT, ADA dQ5 or NL4.3 Vpu. Importantly, the probability of TGN46 overlapping with Vpu, and *vice versa*, was equivalent across the different Vpu variants and mutant (Fig. 8D). Overall, although the data suggest that Vpu downregulates cell-surface HLA-A2 through a mechanism that relies on Vpu redirecting HLA-A2 in a perinuclear compartment that is most probably the TGN, further experimentation is needed to confirm that it is indeed a sequestration.

### Formation of a physical complex between Vpu and HLA-A2 involves the N-terminus/TM domain of Vpu

To further substantiate the confocal data, co-immunoprecipitation experiments were performed to assess whether Vpu and HLA-A2 can form complexes. 293T17 cells were co-transfected with plasmids expressing or not AU1-tagged Vpu and HA-tagged HLA-A*02:01, and subjected to a pulldown with the α-AU1 Ab. As shown in Fig. 9, the ADA WT Vpu consistently formed complexes with HLA-A2 at higher affinity than the NL4.3 Vpu, as evident by the level of co-immunoprecipitated HA-HLA A2 (α-HA, IP blot, Fig. 9A) relative to the amounts of precipitated Vpu (αAU1, IP blot, Fig. 9A). Both ADA dQ5 Vpu and NL4.3 Vpu were significantly less capable of forming the complexes and the NL4.3 was least efficient (Fig. 9B, top panel), thus highlighting the contribution of additional determinants within the N-terminus/TMD of ADA Vpu to the interaction. Indeed, the inclusion of the ADA-NL and NL-ADA Vpu chimeras (Fig. 6) demonstrates that the N-terminus/TMD of the ADA Vpu plays an important role: the ADA-NL Vpu chimera was as efficient as the ADA WT and noticeably better than the NL-ADA (Fig. 9B bottom panel). No HLA-A2 was co-precipitated when cells expressed only Ha-HLA-A2 (lane EV + Ha HLA-A2, Fig. 9A) or Vpu (lane ADA WT, Fig. 9A), validating the specificity of the pull down. Lastly, Vpu did not affect the level of HLA-A2 at steady state (Fig. 9A, Fig. 9C, input blot for Ha), indicating that the downregulation of surface HLA-A2 appears to be governed strictly by a redistribution of the protein. The fact that Vpu did not affect the overall level of HLA-A2 was also observed in infected primary CD4^+^ T cells (Fig. S10). In summary, all three Vpu variants are capable of binding HLA-A2, but doing so with different affinity, suggesting that the ability of Vpu to downregulate cell-surface HLA-A2 is directly related to its capacity to form a high affinity intracellular complex with HLA-A2 through the determinants present in the N-terminus/TMD.

**FIG 9.**
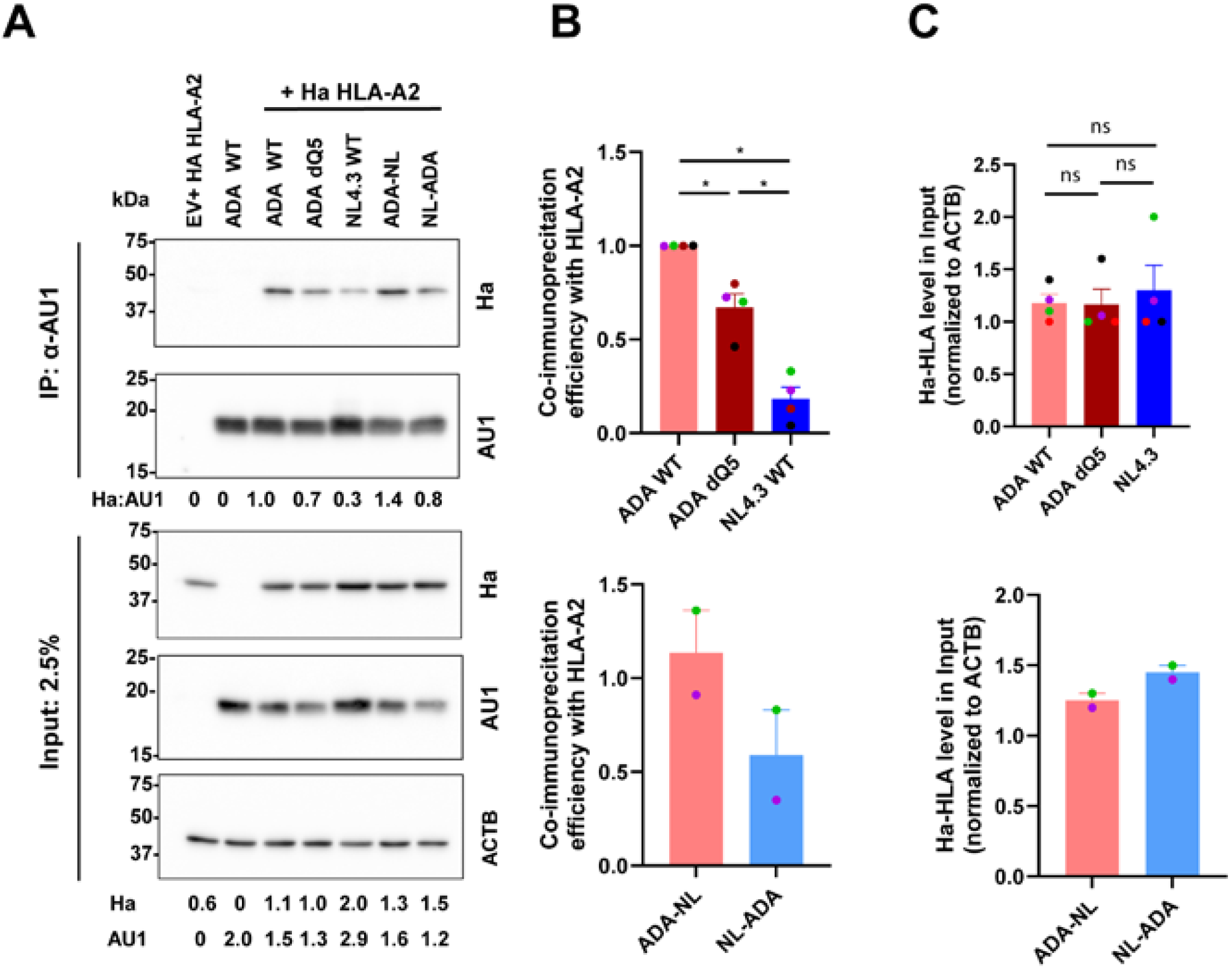
Formation of a physical complex between Vpu and HLA-A2 involves the N-terminus region/TM domain of Vpu. **(A-B)** HEK293T17 were co-transfected with plasmids expressing HA-tagged HLA-A*02:01 and different versions of AU1-tagged Vpu. Co-IP assays were performed using the α-AU1 Ab to pull down the HA-HLA-A2. Panel A: A representative Western blot depicting detection of the indicated proteins in both input lysates and immunoprecipitated fractions. Quantification of protein signals was done using ImageJ, and expression levels of HA and Vpu were determined after normalization to that of actin (ACTB) as shown under the ‘Input’ blots. The numbers shown below the IP blots are the ratio of coprecipitated HLA-A2 normalized to the corresponding precipitated Vpu. The value obtained for ADA WT was set at 1.0. Panels B and C: compiled data (mean +/- SEM) from 2-4 experiments described above. Each dot represents an experiment. When possible, Mann-Whitney’s test was used to test statistical significance. Color-coded experiments clearly demonstrate the importance of the N-terminus/TMD of ADA Vpu in the interactions with HLA-A2. * < 0.05; ns, not significant. See also Fig. S10.

## DISCUSSION

Early studies using the prototypical recombinant strain NL4.3 established the long-standing view that HIV-1 downregulates only HLA-A and HLA-B through Nef (8, 15). Subsequent studies of primary viral strains showed that Vpu and Nef can also modulate HLA-C and HLA-E, respectively (11, 12). Here, we show that Vpu variants from primary strains act independently of Nef to downregulate HLA-A, especially the globally prevalent HLA-A2, and impair TCR-mediated recognition of infected cells. BST2 depletion also reduced HLA-A2-restricted, HIV-specific CD8^+^ T cell signaling. CTL responses were lowest when both Nef and Vpu were present. Vpu proteins from clades A to D of group M and selected groups O and P, but not group N, were sufficient to downregulate HLA-A2. Using ADA Vpu as a reference, we found that Vpu reduces cell-surface HLA-A2 by redirecting the molecule in a perinuclear compartment that colocalizes with TGN46, in a process that involves formation of a physical complex but does lead to degradation of HLA-A2. These findings identify the Vpu N-terminus/transmembrane domain as a key determinant of HLA-A2 downregulation and formation of a high affinity complex.

Vpu contribution to HLA-A modulation was detectable regardless of Nef presence. The data with the WITO virus shown here-in are not dissimilar to earlier observations showing that although Nef played a major role, reduction of cell-surface HLA-A was observable in its absence, suggesting a complementary role by another viral protein (11). The more pronounced contribution of Nef might be related to the timing and its mode of action. Nef is present early in the viral replication cycle and often acts as an adaptor protein to redirect its targets to subcellular compartments for sequestration and/or degradation (reviewed in (47)). In contrast, Vpu is produced later in the replication cycle when structural proteins are expressed (47, 48), and at that stage the pool of HLA-A2 may not be as plentiful, owing to the effect of Nef.

Vpu-mediated modulation of HLA-A expression is conserved in many primary viruses from both the establishment and chronic phases of infection. At least one active Vpu variant was identified in nearly every group M clade, suggesting that pandemic group M viruses are more likely than other groups to reduce HLA-A levels on infected cells. By contrast, this activity was weaker among the less prevalent groups N, O, and P and was undetectable in all tested group N proteins. Analyses of Vpu proteins from ancestral SIVgsn/mus/mon lineages and the SIVcpz MB897 and SIVgor CP2139 precursors suggest that HLA-A modulation arose after SIVgsn/mus/mon contributed to the emergence of Vpu in chimpanzee-and gorilla-infecting SIV lineages, which ultimately gave rise to HIV-1 groups M and O/P, respectively. This evolutionary pattern differs from that of Vpu-mediated HLA-C modulation, which was detected in the ancestral SIVgsn/mus/mon clade (49). Our findings therefore support a model in which Vpu acquired HLA-A downmodulation in chimpanzee SIV, retained it in gorilla SIV and after transmission to humans, the function was conserved to varying degrees across HIV-1 groups and clades. It is tempting to suggest that Vpu complementing Nef-mediated HLA-A downregulation may contribute to infection control and viral persistence. Further analyses of Vpu proteins from additional SIVs, or of HIV-2 Env which mediate some Vpu activities could clarify the evolutionary pathways by which these viruses counteract HLA-A expression or function. Our findings with CH198, CH293, and group N are consistent with previous reports on HLA-C (11, 42, 43). Group N Vpu proteins may have acquired BST2 antagonism at the expense of activities against other host factors, including CD4, NTB-A, CD1d, and HLA-I (43, 44).

The extent of HLA-A2 modulation varies among primary Vpu variants, with some showing modest or undetectable effects. Because Vpu-mediated downregulation of HLA-A and BST2 depletion independently impairs TCR recognition of infected cells, strong BST2 antagonism by Vpu may reduce selective pressure for Vpu-mediated HLA-A downregulation. Alternatively, variation among Vpu proteins may reflect differences in HLA-A allele expression or the strength of CTL-mediated immune pressure. Indeed, distinct HLA-C alleles impose different selective pressures on HIV-1, with higher HLA-C expression associated with stronger downregulation (10). Similarly, when antigen presentation by a given HLA molecule elicits a weak CTL response, the virus may be less likely to evolve mechanisms that evade immune recognition. This possibility is illustrated by the prototypical NL4.3 strain, which was generated from viruses that had been extensively propagated *in vitro* without immune pressure and has lost Vpu activity against both HLA-C and HLA-A.

HIV-1 Nef has been proposed to reduce cell-surface HLA-A2 through several mechanisms, including impaired transport (45), degradation (7), and sequestration (26). In contrast, our data suggest that Vpu primarily retains HLA-A2 intracellularly, as it does other non-degraded targets such as CD1d (46) and NTB-A (47). Vpu interacts with HLA-A2 through its N-terminus/TMD, and the strength or stability of this interaction may determine its ability to redirect HLA-A2 in a PNC and reduce its surface expression. Consistent with this model, glutamine at position 5 is required for ADA Vpu to interact with HLA-A2 and redirect it, thereby reducing its cell-surface expression. Subtype B Vpu proteins that strongly downregulated HLA-A2 often contained glutamine or glutamic acid at this position, whereas less active or inactive variants generally lacked these residues. Isoleucine at position 6 of ADA Vpu was also important: replacing it with threonine, the corresponding residue from the weakly active 2-87 Vpu, reduced HLA-A2 downregulation, further supporting the importance of specific residues in this region.

While a previous report (10) did not detect an interaction of Vpu with HLA-A, it is worth mentioning that the co-immunoprecipitation experiments used in the other study were conducted with Vpu from primary strain 2_87, which were found in both studies to modestly downregulate HLA-A2. More significantly, there were also differences in the experimental systems regarding stringency of the lysis/binding buffers (detergent composition) and the type and position of the tag in Vpu (Flag in the other paper *vs.* AU in this study; N-terminus in the other paper *vs.* C-terminus in this paper) that could have contributed to the better detection of the binding in our study. Delineating precisely the positions within Vpu and HLA-A that are critical for the interactions and assessing whether they overlap with those required for HLA-C downregulation and binding would be of interest and require further investigation.

Although HLA-A2 presents significantly fewer viral peptides than other HLA-I molecules, it has the highest presentation of Gag-derived peptides to CD8^+^ T cells (49), a feature that is often associated with immune protection in HIV infection. With Vpu complementing Nef activity, HIV-1 ensures that antigen presentation is efficiently reduced and effector functions by T cells evaded late in the infection cycle when viral production is at its peak and immune evasion is most critical. The fact that BST2 depletion was associated with reduced HIV-1-specific TCR-mediated signaling points to another mechanism whereby Vpu could evade immune responses by limiting antigen processing and presentation to CD8^+^ T cells. Taken together, this study highlights functions of Vpu that are less appreciated owing to a high level of functional redundancy between Nef and Vpu. Our findings reveal how HIV has evolved to use multiple proteins to avoid immune recognition and ultimately, immune evasion. In the context of the HIV reservoir persisting during antiretroviral therapy (ART), CTLs are important in controlling and limiting the size of the reservoir. The fact that both Nef and Vpu can downregulate HLA-I and limit viral antigen presentation indicates that even if latently infected cells are sufficiently reactivated by latency reversing agents, the viral reservoir might not be efficiently cleared by CTLs. This is yet another hurdle that cure research must overcome.

## LIMITATION OF STUDY

While HIV-1-induced HLA-A2 modulation was well correlated with the impairment of TCR-mediated signaling in the reporter cell system, the findings were not fully recapitulated in a CTL response assay using primary CD8^+^ T cells. In addition, the finding that Vpu redirects HLA-A2 to a PNC was done in HeLa cells since changes in protein subcellular localization are easier to visualize in epithelial cells than in T cells with narrow cytoplasm. Lastly, the evidence of high affinity complex formed between ADA Vpu and HLA-A2 was obtained in an overexpression system. Replicating these results in infected CD4^+^ T cells would further validate the interactions.

## ACKNOWLEDGEMENTS

The following reagents were obtained through the NIH AIDS Reagent Program, Division of AIDS, NIAID, and NIH: Panel of full-length transmitted/founder (T/F) HIV-1 Infectious Molecular Clones, ARP-8129, contributed by Dr. John Kappes. We thank Dr. Scott G. Kitchen (UCLA) for providing the pCCL.PPT.hPGK.1.9.IRES.eGFP vector. We are grateful for the technical support provided by the IRCM cytometry platform. We thank all the blood donors and the IRCM clinic staff for their valuable contributions. This study was supported by the Canadian Institutes of Health Research (CIHR) grants FDN-154324 and HB2-164064 (Canadian HIV Cure Enterprise-CanCURE) to ÉAC. This work was also supported in part by the Fonds de Recherche du Québec-Santé (FRQS) supported Réseau SIDA/Maladies Infectieuses. ÉAC is the recipient of the IRCM-Université de Montréal Chair of Excellence in HIV Research. The funders had no role in study design, data collection and interpretation, or the decision to submit the work for publication.

## AUTHOR CONTRIBUTIONS

Conceptualization, M.G.B, T.N.Q.P., E.A.C; Methodology, M.G.B., T.N.Q.P., C.C., M.A.B, F.K; Investigation, M.G.B., T.N.Q.P., C.C., V.S., M.L., F.D., W.M. Resources, M.A.B., F.K., E.A.C; Writing, M.G.B, T.N.Q.P. and E.A.C; Funding Acquisition and Supervision, E.A.C.

## MATERIALS AND METHODS

### Cell lines

The human embryonic kidney (HEK) 293T cell line (RRID:CVCL_0063; female cells) and HeLa (RRID:CVCL_0030; female cells) were obtained from the American Type Culture Collection (ATCC). TZM-bl (RRID: CVCL_B478), derived from HeLa cells (female origin), were acquired through the NIH AIDS Reagent Program. The HeLa HA-tagged A*02:01 cell line was generated in house as described below. CEM-A2 T cell line was generated from CEM-SS line (50). CEM CD4^+^ T cells (RRID: CVCL_0207) were of female origin. CEM-CCR5 cells were derived from CEM CD4^+^ T (51). Jurkat E6.1 CD4^+^ T cells were of a male. PT67 packaging cell lines (RRID:CVCL_9093) were from the ATCC.

BST2-depleted CEM-A2 cell line was made using lentiviral vectors expressing BST2-targeting shRNA (Sh45) or scramble control. Lentiviral particles were made as described below.

HeLa HA-tagged A*02:01 cell line was generated using retroviral particles expressing HLA-A*02:01. In brief, an HA tag was inserted by overlapping PCR after the leader peptide sequence of HLA-A*0201 in pMSCV-A*02:01 (50). PT67 cells were transfected with the modified pMSCV-A*02:01 and SVCMV-IN VSVg (52) to produce retroviral particles. Supernatants were purified over a 20% sucrose cushion 48 h later, and the particles were used to generate the HeLa HA-tagged A*02:01 line (HeLa-A2).

HEK293T, HeLa-based and PT67 cell lines were cultivated in DMEM media. CEM-based and Jurkat T cells were grown in RPMI-1640. All growth media were supplemented with 10% fetal bovine serum (FBS). Cells were maintained in incubators supplied with 5% CO_2_ in air atmosphere (37^0^C). All cell lines were routinely checked for mycoplasma. No other information related to cell authentication was available.

### Primary cell cultures from human subjects

Peripheral blood samples were obtained from healthy adult donors who gave written informed consent in accordance with the Declaration of Helsinki under research protocols approved by the IRCM Human Research Ethics Review Board (IRCM 2012-16). PBMCs were isolated by Ficoll-Paque centrifugation (GE Healthcare) and cultured in RPMI-1640 (+ 10% FBS). CD4^+^ and CD8^+^ T lymphocytes were isolated using a CD4^+^ T Cell Isolation Kit and CD8 microbeads, respectively (both were from Miltenyi Biotec). Enriched CD4^+^ T cells were activated with PHA-L (5 µg/mL) for 48 h and infected with HIV 5 days post-isolation. Purified CD8^+^ T cells were activated with immobilized α-CD3 Ab and soluble α-CD28 Ab (both were from BioLegend and used at 2 µg/ml) for 24h, washed and transduced with the lentiviral vector expressing the HLA-A*02:01-restricted, HIV Gag epitope (SL9)-specific TCR for 3-5 days before being put in co-culture with infected CEM-A2 cells.

### Plasmids and cell transfection

Bicistronic CMV-promoter-based pCGCG vectors co-expressing Vpu and the enhanced green fluorescent protein (eGFP) were described previously (53). The origin and source of the virus from where the Vpu variants were cloned into the pCGCG vectors were as detailed previously (53). In brief, Vpu variants were derived from an infectious molecular clone of the founder virus (ZM246F and ZM247F) or chemically synthesized from viral sequences that were originally amplified from blood cells of patients with chronic HIV infection (SE.95, CY.94, KE.00, UG.99, ZA.90 and CM.02). Similarly, the SIV Vpus from EK505, MB897 and CP2139 were obtained from either fecal viral RNA or from blood. The Vpu variants were not codon optimized. The AU1 tag along with XbaI and MluI restriction sites flanking the reading frames were introduced in the pCGCG vector by PCR. The AU1 tag was fused to the C-terminus of Vpu and had been shown to not affect Vpu functions in promoting viral particle release through BST2 antagonism or CD4 downregulation (53).

HeLa-A2 cells were transfected with a GFP-marked plasmid expressing EV or AU1-tagged Vpu (0.5 to 2 μg of total DNA) using Lipofectamine 2000 or polyethylenimine (PEI). Empty plasmid pcDNA3.1 was used to equilibrate the amounts of transfected DNA. Cells were processed for flow cytometry or Western blotting at 24 h or 48 h post transfection depending on experiments.

### HIV proviral constructs and production of lentiviral particles

All GFP-marked viruses used were derived from a pNL4.3 construct, in which the *nef* gene is followed by an internal ribosome entry site that allows for GFP expression (pNL4.3-GFP) (8). Vpu-deficient mutant (dU) was generated by subcloning a SalI-KpnI fragment from the Vpu-defective pNLVpuDEL1 (54) into the pNL4.3-GFP. CCR5-tropic GFP-marked proviral DNA (pNL4.3-ADA-GFP) and Vpu-deficient mutant were reported previously (55). In brief, the pNL4.3-ADA-GFP WT proviral construct was generated by transferring, the 2.7-kb SalI-BamHI fragment of the NLHXADA into the pNL4.3 backbone at these two sites. The SalI-BamHI cassette contains a SacI-BsmI insert encoding for

ADA Vpu and Env gp120 and 2 flanking sequences SalI-SacI and BsmI-BamHI of HxB2 which encode for the first and second exons of Tat and Rev. The insertion of this DNA fragment rendered NL4.3 macrophage-tropic (56) through the ability of its recombinant Env containing the ADA determinant to mediate viral entry via a CCR5-dependent mechanism (57). WT and dU NL4.3 viruses were rendered defective for Nef gene by a frameshift using XhoI digest and refilling DNA using T4 DNA polymerase. The full-length T/F infectious molecular clone WITO virus was obtained through the NIH AIDS Reagent Program (58–61). Vpu-deficient WITO was generated by subcloning a PCR-generated EcoRI-BamH1 Vpu-defective fragment (34). To generate Nef-deficient WITO, the additional XhoI restriction site in the backbone multiple cloning site (MCS) was removed by excising the NotI-XbaI fragment and refilling the DNA using T4 DNA polymerase. MCS-deleted WT and dU WITO viruses were rendered defective for the *nef* gene as described above for NL4.3 viruses.

Viruses were pseudotyped by transfecting HEK293T cells with 25 µg proviral DNA and 8 µg psvCMV-VSV-G using Lipofectamine 2000 or calcium phosphate-based method. To make lentiviral vectors expressing shRNA targeting BST2 or scramble control, 293T cells were co-transfected with puromycin-resistant plasmid pLKO.1 expressing shRNA targeting BST2 (Clone ID: TRCN0000107018, OpenBiosystems; Sh45) or scramble control along with packaging plasmid psPAX2 (a gift from Dr. Trono, Swiss Federal Institute of Technology) and psvCMV-VSV-G. Alternatively, to produce vectors expressing the GFP-marked HLA-A2 restricted, SL9-specific TCR, 293T cells were co-transfected with the pCCL.PPT.hGPK.1.9.IRES.eGFP (a kind gift from Dr. Scott G. Kitchen, UCLA) (62) along with the psPAX2 and psvCMV-VSV-G. Viral supernatant was harvested between 48-60 h post transfection and purified over a 20% sucrose gradient (33). Virus titration was performed in CEM.CCR5 CD4^+^ T cells. Depending on viral titers, different volumes of viruses maybe used to achieve comparable infection rates in target cells.

### HIV infection, co-cultures of CD4^+^ T cells with CD8^+^ T cells and analysis

Activated CD4^+^ T cells and CEM-A2 were infected at multiplicity of infection (MOI) of 1 for up to 72 h. In the experiment aimed at examining transcriptional levels of *HLA-A*02:01* gene, infected cells were sorted using flow cytometry. GFP^+^ cells (in the case of the NL4.3 and ADA viruses) and gp120^+^ cells (in the case of WITO) along with the respective uninfected (mock) cells were extracted for total RNA and analyzed by RT-qPCR.

The experiment aimed to assess TCR-mediated signalling was carried out as follows. At 48 h post infection, CEM-A2 cells were analyzed by flow cytometry for infection rates (normally around 50-70% for each virus) and the same number of infected CEM-A2 T cells (250,000 GFP^+^ or p24^+^ cells) were co-cultured for up to 24 h with reporter Jurkat T cells which co-express CD8 and a TCR specific for the HLA-A*02:01-restricted HIV gag epitope FK10 (FLGKIWPSYK; residues 433 to 442) (50). The effector : target ratios varied between 1 to 2. In brief, Jurkat E6.1 cells (10^7^) resuspended at 2.5 × 10^7^ cells/mL in Opti-MEM were transferred to 0.4 cm cuvettes (Bio-Rad Laboratories, Inc.) containing 3 μg pSELECT-5B2alpha, 3 μg pSELECT-5B2beta, 5 μg pORF9-hCD8A and 10 μg pNFAT-luciferase, incubated for 10 min at room temperature (RT) and transfected using a Gene Pulser Xcell™ Electroporation System (Bio-Rad Laboratories, Inc.) with the following setting: 3 square-wave protocol; 350 V; 5 millisecond pulse length and 2 pulses with 3 sec gap as pulse interval. After a 10 min recovery at RT, cells were incubated for 16–20 h at 37°C in 5 mL RPMI containing 10% FBS and penicillin/streptomycin. Prior to co-culture with CEM-A*02:01 cells, flow cytometry analysis was performed to confirm CD8 and TCR expression on Jurkat cells. Infection rates were determined on the day of the co-cultures, and the same number of target cells was co-cultured with effector cells. CD8 presence was validated using an α-CD8 Ab and TCR presence by GFP expression. After the co-culture, cells were lysed and analyzed for luciferase expression using bioluminescence.

The experiment aimed to assess HIV-specific CTL responses were performed as follows. CD8^+^ T cells expressing a GFP-marked, HLA-A*02:01-restricted TCR specific for the, HIV Gag SL9 epitope (SL9) TCR (62) were co-cultured for 3 h with CEM-A2 cells that had been infected for 24 h with WITO viruses for 24 h. Both target and effector cells were washed thoroughly with PBS before being placed together. To normalize target cell input, cultures were supplemented with uninfected cells so that each condition contained the same number of p24^+^ target cells at the start of the co-culture (5,000 target cells per condition). The frequency of GFP^+^ CD8^+^ T cells was determined by flow cytometry before the co-culture, and the effective SL9-specific effector : p24^+^ target ratio ranged from 1:2 to 1:5. Monensin (BD Biosciences) and PE-conjugated α-CD107a Ab (BioLegend) were present throughout the 3-h co-culture. Brefeldin A (BioLegend) was added during the final 2 h. There-after, spent media was removed and cells were washed with PBS before being successively stained with live/dead cell exclusion dye Zombie Aqua (BioLegend) and AF647-conjugated α-CD8 Ab (BioLegend). Cells were fixed and permeabilized using Cytofix/Cytoperm kit (BD Biosciences) and stained for IFNγ and TNFα (PE-Cy7-conjugated α-IFNγ Ab and Pe-dazzle-conjugated α-TNFα Ab were from BioLegend). GFP-marked CTLs were then analyzed by flow cytometry for cell surface CD107a (indicative of cell degranulation) and cytokine expression.

### Flow cytometry-based assays

Cells were washed with cold PBS/EDTA/FBS, blocked with human IgG when applicable, and stained with purified α-HLA-A2 (BioLegend, clone BB7.2), AF647-conjugated α-HLA-A2 (BioLegend, clone BB7.2), PE-conjugated α-HLA-A (BD BioSciences, clone 1082C5) or unconjugated α-HLA-C (BioLegend, clone DT9) Abs for 60 min at 4°C. Cells stained with relevant isotype controls served as negative controls. Relevant secondary Abs were used as appropriate to reveal the fluorescence signals of HLA-A2 or HLA-C. Following surface staining, cells were fixed and permeabilized using Cytofix/Cytoperm kit (BD Biosciences) and stained for HIV-1 p24 or AU1-Vpu using mouse α-HIV core antigen Ab (Beckman Coulter, KC57-FITC) or rabbit α-AU1 Ab (LSBio, cat# LS-C83247-100), respectively. For the detection and sorting of WITO-infected cells for RT-qPCR analysis, target cells were stained for gp120 using the human 10-1074 Ab (NIH AIDS Reference Reagents). Cells were incubated for 30 min at room temperature, followed by another 30 min at 4°C. In all relevant cases, cell incubation with appropriate secondary Abs was done at 4°C for 30-45 min.

To assess for the rate of HLA-A2 internalization, HeLa-A2 cells were labelled on ice with unconjugated mouse α-human HLA-A2 Ab (BioLegend, clone BB7.2) for 90 min 12h post transfection. Cells were washed and chased for 4 h at 37°C. At indicated time points starting 30 min thereafter, cells were surface stained with AF674-conjugated α-mouse secondary Ab (Thermo Fisher Scientific, cat # A-31571) and analyzed by flow cytometry for HLA-A2 on the transfected (GFP^+^) cells. Staining at time 0 was performed to gauge the protein level at the start of the chase.

Samples were acquired on a BD LSRFortessa™ Cell Analyzer (BD Bioscience) using FACSDiva software or on a Cyan ADP cytometer (Beckman Coulter) and analyzed by FlowJo (Versions 9.9.3 and 10.1).

### Coimmunoprecipitation

293T17 plated in 6-well plates the day before were transfected for 48 h with 500 ng pCGCG EV or 500-1750 ng constructs expressing different versions of Vpu. When relevant, 293T cells were co-transfected with 250 ng pSELECT Ha-HLA-A2. PcDNA3.1 EV was used to equilibrate the total amounts of DNA transfected (2 μg). Cells were lysed by resuspending the cell pellet in 200 µl RIPA-DOC buffer (1% NP-40, 140 mM NaCl, 0.5% SDS, 1.2 mM Na-Deoxycholate, 8 mM Na2HPO4 and 2 mM NaH2PO4, protease inhibitor mix, pH 7.2). The lysates were immediately diluted in 400 µl of the RIPA-based buffer (RIPA-DOC buffer without the detergents), pipetting several times before a nucleus burst. Samples were incubated on ice for 30 min, and clarified at 15,000 *x g* for 20 min at 4°C. The lysates were precleared for 30 min at 4°C in 12.5 µl Dynabeads protein G and then placed on a magnet. During the preclearing step, α-AU1 Ab (3.5 µg) was complexed to Dynabeads protein G (25 µl in 100 µl PBS, Thermo Fisher Scientific, cat# 10003D) for 30 min at RT (on a rotator). Immunoprecipitation was carried out with 800-1000 µg precleared protein lysates for 3 h at 4°C (on rotation). The beads were washed three times in 500 µl IP buffer (RIPA-based containing 0.167% SDS and 0.4 mM Na-Deoxycholate) and once in PBS + 1% Triton. Immunocomplexes were eluted by boiling in Laemmli buffer, electrophoresed by SDS-PAGE and processed for Western blotting.

### Western Blotting

Cells were washed with phosphate-buffered saline (PBS) and then lysed in RIPA-DOC buffer for 20 min at 4°C. Immunoblotting was performed according to standard protocols. Samples were incubated for overnight at 4°C with either the α-Ha (BioLegend, clone 16B12), α-AU1, α-ACTB (α-actin, MP Biomedicals, cat# SKU08691001), or α-GFP (Sigma-Aldrich, cat# SAB4301138) Abs. Unbound Abs were washed extensively and probed with the corresponding secondary Ab conjugated with horseradish peroxidase for 1 h.

### Immunofluorescence

HeLa-A2 cells were transfected for 24 h with EV plasmid, or with constructs expressing AU1-tagged NL4.3, ADA WT or ADA dQ5 Vpu. Cells were washed with PBS containing 1 mM MgCl_2_, fixed in 3.7% formaldehyde and permeabilized with 0.5% Triton X-100/PBS for 20 min. Immunodetection of AU1-Vpu, HA-HLA-A2 and TGN (TGN46) was performed using the rabbit α-AU1, mouse α-HA and sheep α-TGN46 Ab (GenTex, cat# GTX74290), respectively. AlexaFluor-488 conjugated α-rabbit, AlexaFluor-647 conjugated α-mouse and AlexaFluor-555 conjugated α-sheep Abs were all from ThermoFisher Scientific and used to visualize AU1, Ha and TGN46, respectively. Hoechst 33342 (Sigma Aldrich, cat# B2261) stained cell nuclei. Cells were examined by confocal microscopy, using a LSM 710 imaging system mounted on a Zeiss AxioObserver Z1 inverted microscope equipped with a Plan-Apochromat 63x/1.40 Oil DIC M27 objective and controlled by Zen Black 2009 v6.0 (Carl Zeiss, Jena, Germany).

#### Co-distribution Analysis

Individual pixel analysis for co-distribution of Vpu (green fluorescence signal) in Golgi or with HLA A2 (red fluorescence signal) was performed using the “colocalization threshold” plugin in ImageJ. Confocal image stacks were cleared of the background noise using the mean intensity calculated from the region of interest (ROI) outside the cells and the “subtract’’ function from the process menu, resulting in an image filtered to remove point noise. The single channel-specific threshold-adjusted Mander’s coefficient (tM) provides a good indicator of the proportion of the green signal (Vpu) coinciding with the signal in the red channel (Golgi or HLA A2). ImageJ 1.54n (National Institute of Health) was used to process the entire image before cropping to emphasize the main point of the image. Processing was limited to background subtraction and brightness/contrast adjustments.

### Nucleic-acid extraction and RT-qPCR

Total RNA was extracted using QIAzol Lysis Reagent (QIAGEN Sciences, USA) according to manufacturer’s instructions. Total RNA (300 ng) was reverse transcribed using SuperScript IV RT (Thermo Fisher Scientific, U.S.A). SYBR-based PCR was done using primers: BST2: 5’-CAGAAGGGCTTTCAGGATG-3’ (forward); 5’-TTTGTCCTTGGGCCTTCTC -3’ (reverse); GAPDH: 5’-GCCATCAATGACCCCTTCATT-3’ (forward); 5’-TTGACGGTGCCATGGAATTT-3’ (reverse). TaqMan-based real-time PCR was done using primers: HLA-A*02:01: 5’-GGACGGGGAGACACGGAAAG-3’ (forward); 5’-ACACCGTCCAGAGGATGTATGGC-3’ (reverse); 5’-GACTCACCGAGTGG-3’ (probe; dye FAM; quencher ZEN-3’ Iowa Black FQ).

## Quantification and statistical analysis

Quantification of Western blots was performed using ImageJ software. Data analysis including statistics was done using GraphPad Prism (Version 8.0). Descriptive measures (mean, median, and percent) were used to present graphically the compiled data. Nonparametric Mann-Whitney’s U-test was conducted to compare ranks between two groups. When comparing multiple groups, non-parametric tests Kruskal-Wallis or Friedman test were performed, followed by Dunn’s multiple comparison. Correlation analysis was performed using a two-tailed Spearman test. A *p* value of less than 0.05 was considered statistically significant.

## Data Availability

The data will be made fully available and without restriction.

## Supplemental Material

Supplemental Figures. Figures S1-S10.

## REFERENCES

1. Martin MP, Carrington M. 2013. Immunogenetics of HIV disease. Immunol Rev 254:245–64.

2. Langford SE, Ananworanich J, Cooper DA. 2007. Predictors of disease progression in HIV infection: a review. AIDS Res Ther 4:11.

3. Singh KK, Spector SA. 2009. Host genetic determinants of human immunodeficiency virus infection and disease progression in children. Pediatr Res 65:55R–63R.

4. McLaren PJ, Fellay J. 2021. HIV-1 and human genetic variation. Nat Rev Genet 22:645–657.

5. Crux NB, Elahi S. 2017. Human Leukocyte Antigen (HLA) and Immune Regulation: How Do Classical and Non-Classical HLA Alleles Modulate Immune Response to Human Immunodeficiency Virus and Hepatitis C Virus Infections? Front Immunol 8:832.

6. Beltrami S, Rizzo S, Strazzabosco G, Gentili V, Alogna A, Narducci M, Bortolotti D, Schiuma G, Rizzo R. 2023. Non-classical HLA class I molecules and their potential role in viral infections. Hum Immunol 84:384–392.

7. Roeth JF, Williams M, Kasper MR, Filzen TM, Collins KL. 2004. HIV-1 Nef disrupts MHC-I trafficking by recruiting AP-1 to the MHC-I cytoplasmic tail. J Cell Biol 167:903–13.

8. Cohen GB, Gandhi RT, Davis DM, Mandelboim O, Chen BK, Strominger JL, Baltimore D. 1999. The selective downregulation of class I major histocompatibility complex proteins by HIV-1 protects HIV-infected cells from NK cells. Immunity 10:661–71.

9. Lanier LL. 2005. NK cell recognition. Annu Rev Immunol 23:225–74.

10. Bachtel ND, Umviligihozo G, Pickering S, Mota TM, Liang H, Del Prete GQ, Chatterjee P, Lee GQ, Thomas R, Brockman MA, Neil S, Carrington M, Bwana B, Bangsberg DR, Martin JN, Kallas EG, Donini CS, Cerqueira NB, O’Doherty UT, Hahn BH, Jones RB, Brumme ZL, Nixon DF, Apps R. 2018. HLA-C downregulation by HIV-1 adapts to host HLA genotype. PLoS Pathog 14:e1007257.

11. Apps R, Del Prete GQ, Chatterjee P, Lara A, Brumme ZL, Brockman MA, Neil S, Pickering S, Schneider DK, Piechocka-Trocha A, Walker BD, Thomas R, Shaw GM, Hahn BH, Keele BF, Lifson JD, Carrington M. 2016. HIV-1 Vpu Mediates HLA-C Downregulation. Cell Host Microbe 19:686–95.

12. van Stigt Thans T, Akko JI, Niehrs A, Garcia-Beltran WF, Richert L, Sturzel CM, Ford CT, Li H, Ochsenbauer C, Kappes JC, Hahn BH, Kirchhoff F, Martrus G, Sauter D, Altfeld M, Holzemer A. 2019. Primary HIV-1 Strains Use Nef To Downmodulate HLA-E Surface Expression. J Virol 93.

13. Adachi A, Gendelman HE, Koenig S, Folks T, Willey R, Rabson A, Martin MA. 1986. Production of acquired immunodeficiency syndrome-associated retrovirus in human and nonhuman cells transfected with an infectious molecular clone. J Virol 59:284–91.

14. Schwartz O, Marechal V, Le Gall S, Lemonnier F, Heard JM. 1996. Endocytosis of major histocompatibility complex class I molecules is induced by the HIV-1 Nef protein. Nat Med 2:338–42.

15. Collins KL, Chen BK, Kalams SA, Walker BD, Baltimore D. 1998. HIV-1 Nef protein protects infected primary cells against killing by cytotoxic T lymphocytes. Nature 391:397–401.

16. Specht A, Telenti A, Martinez R, Fellay J, Bailes E, Evans DT, Carrington M, Hahn BH, Goldstein DB, Kirchhoff F. 2010. Counteraction of HLA-C-mediated immune control of HIV-1 by Nef. J Virol 84:7300–11.

17. Sugden SM, Bego MG, Pham TN, Cohen EA. 2016. Remodeling of the Host Cell Plasma Membrane by HIV-1 Nef and Vpu: A Strategy to Ensure Viral Fitness and Persistence. Viruses 8:67.

18. Ramirez PW, Sharma S, Singh R, Stoneham CA, Vollbrecht T, Guatelli J. 2019. Plasma Membrane-Associated Restriction Factors and Their Counteraction by HIV-1 Accessory Proteins. Cells 8.

19. Barone ME, Lim A, Woody M, Taklifi P, Yeasmin F, Wang K, Lewinski MK, Singh R, Stoneham CA, Jia X, Guatelli J. 2025. Adaptor Protein Complexes in HIV-1 Pathogenesis: Mechanisms and Therapeutic Potential. Viruses 17.

20. Tokarev A, Guatelli J. 2011. Misdirection of membrane trafficking by HIV-1 Vpu and Nef: Keys to viral virulence and persistence. Cell Logist 1:90–102.

21. Kirchhoff F. 2010. Immune evasion and counteraction of restriction factors by HIV-1 and other primate lentiviruses. Cell Host Microbe 8:55–67.

22. Jia B, Serra-Moreno R, Neidermyer W, Rahmberg A, Mackey J, Fofana IB, Johnson WE, Westmoreland S, Evans DT. 2009. Species-specific activity of SIV Nef and HIV-1 Vpu in overcoming restriction by tetherin/BST2. PLoS Pathog 5:e1000429.

23. Zhang F, Wilson SJ, Landford WC, Virgen B, Gregory D, Johnson MC, Munch J, Kirchhoff F, Bieniasz PD, Hatziioannou T. 2009. Nef proteins from simian immunodeficiency viruses are tetherin antagonists. Cell Host Microbe 6:54–67.

24. Douglas JL, Viswanathan K, McCarroll MN, Gustin JK, Fruh K, Moses AV. 2009. Vpu directs the degradation of the human immunodeficiency virus restriction factor BST-2/Tetherin via a betaTrCP-dependent mechanism. J Virol 83:7931–47.

25. Mitchell RS, Katsura C, Skasko MA, Fitzpatrick K, Lau D, Ruiz A, Stephens EB, Margottin-Goguet F, Benarous R, Guatelli JC. 2009. Vpu antagonizes BST-2-mediated restriction of HIV-1 release via beta-TrCP and endo-lysosomal trafficking. PLoS Pathog 5:e1000450.

26. Neil SJ, Zang T, Bieniasz PD. 2008. Tetherin inhibits retrovirus release and is antagonized by HIV-1 Vpu. Nature 451:425–30.

27. Serra-Moreno R, Zimmermann K, Stern LJ, Evans DT. 2013. Tetherin/BST-2 antagonism by Nef depends on a direct physical interaction between Nef and tetherin, and on clathrin-mediated endocytosis. PLoS Pathog 9:e1003487.

28. Perez-Caballero D, Zang T, Ebrahimi A, McNatt MW, Gregory DA, Johnson MC, Bieniasz PD. 2009. Tetherin inhibits HIV-1 release by directly tethering virions to cells. Cell 139:499–511.

29. Preusser A, Briese L, Baur AS, Willbold D. 2001. Direct in vitro binding of full-length human immunodeficiency virus type 1 Nef protein to CD4 cytoplasmic domain. J Virol 75:3960–4.

30. Grzesiek S, Stahl SJ, Wingfield PT, Bax A. 1996. The CD4 determinant for downregulation by HIV-1 Nef directly binds to Nef. Mapping of the Nef binding surface by NMR. Biochemistry 35:10256–61.

31. Cluet D, Bertsch C, Beyer C, Gloeckler L, Erhardt M, Gut JP, Galzi JL, Aubertin AM. 2005. Detection of human immunodeficiency virus type 1 Nef and CD4 physical interaction in living human cells by using bioluminescence resonance energy transfer. J Virol 79:8629–36.

32. Williams M, Roeth JF, Kasper MR, Fleis RI, Przybycin CG, Collins KL. 2002. Direct binding of human immunodeficiency virus type 1 Nef to the major histocompatibility complex class I (MHC-I) cytoplasmic tail disrupts MHC-I trafficking. J Virol 76:12173–84.

33. Dube M, Roy BB, Guiot-Guillain P, Binette J, Mercier J, Chiasson A, Cohen EA. 2010. Antagonism of tetherin restriction of HIV-1 release by Vpu involves binding and sequestration of the restriction factor in a perinuclear compartment. PLoS Pathog 6:e1000856.

34. Cong L, Sugden SM, Leclair P, Lim CJ, Pham TNQ, Cohen EA. 2021. HIV-1 Vpu Promotes Phagocytosis of Infected CD4(+) T Cells by Macrophages through Downregulation of CD47. mBio 12:e0192021.

35. Dube M, Paquay C, Roy BB, Bego MG, Mercier J, Cohen EA. 2011. HIV-1 Vpu antagonizes BST-2 by interfering mainly with the trafficking of newly synthesized BST-2 to the cell surface. Traffic 12:1714–29.

36. Arrieta-Bolanos E, Hernandez-Zaragoza DI, Barquera R. 2023. An HLA map of the world: A comparison of HLA frequencies in 200 worldwide populations reveals diverse patterns for class I and class II. Front Genet 14:866407.

37. Nomaguchi M, Fujita M, Adachi A. 2008. Role of HIV-1 Vpu protein for virus spread and pathogenesis. Microbes Infect 10:960–7.

38. Basmaciogullari S, Pizzato M. 2014. The activity of Nef on HIV-1 infectivity. Front Microbiol 5:232.

39. Sauter D, Hotter D, Van Driessche B, Sturzel CM, Kluge SF, Wildum S, Yu H, Baumann B, Wirth T, Plantier JC, Leoz M, Hahn BH, Van Lint C, Kirchhoff F. 2015. Differential regulation of NF-kappaB-mediated proviral and antiviral host gene expression by primate lentiviral Nef and Vpu proteins. Cell Rep 10:586–99.

40. Langer S, Hammer C, Hopfensperger K, Klein L, Hotter D, De Jesus PD, Herbert KM, Pache L, Smith N, van der Merwe JA, Chanda SK, Fellay J, Kirchhoff F, Sauter D. 2019. HIV-1 Vpu is a potent transcriptional suppressor of NF-kappaB-elicited antiviral immune responses. Elife 8.

41. Boegel S, Lower M, Bukur T, Sahin U, Castle JC. 2014. A catalog of HLA type, HLA expression, and neo-epitope candidates in human cancer cell lines. Oncoimmunology 3:e954893.

42. Bibollet-Ruche F, Heigele A, Keele BF, Easlick JL, Decker JM, Takehisa J, Learn G, Sharp PM, Hahn BH, Kirchhoff F. 2012. Efficient SIVcpz replication in human lymphoid tissue requires viral matrix protein adaptation. J Clin Invest 122:1644–52.

43. Raehtz K, Pandrea I, Apetrei C. 2016. The well-tempered SIV infection: Pathogenesis of SIV infection in natural hosts in the wild, with emphasis on virus transmission and early events post-infection that may contribute to protection from disease progression. Infect Genet Evol 46:308–323.

44. Bell SM, Bedford T. 2017. Modern-day SIV viral diversity generated by extensive recombination and cross-species transmission. PLoS Pathog 13:e1006466.

45. Takehisa J, Kraus MH, Ayouba A, Bailes E, Van Heuverswyn F, Decker JM, Li Y, Rudicell RS, Learn GH, Neel C, Ngole EM, Shaw GM, Peeters M, Sharp PM, Hahn BH. 2009. Origin and biology of simian immunodeficiency virus in wild-living western gorillas. J Virol 83:1635–48.

46. Hauser H, Lopez LA, Yang SJ, Oldenburg JE, Exline CM, Guatelli JC, Cannon PM. 2010. HIV-1 Vpu and HIV-2 Env counteract BST-2/tetherin by sequestration in a perinuclear compartment. Retrovirology 7:51.

47. Kirchhoff F, Schindler M, Specht A, Arhel N, Munch J. 2008. Role of Nef in primate lentiviral immunopathogenesis. Cell Mol Life Sci 65:2621–36.

48. Schwartz S, Felber BK, Fenyo EM, Pavlakis GN. 1990. Env and Vpu proteins of human immunodeficiency virus type 1 are produced from multiple bicistronic mRNAs. J Virol 64:5448–56.

49. Dodd GK, Kesmir C. 2025. In silico functional analysis of the human, chimpanzee, and gorilla MHC-A repertoires. Immunogenetics 77:12.

50. Anmole G, Kuang XT, Toyoda M, Martin E, Shahid A, Le AQ, Markle T, Baraki B, Jones RB, Ostrowski MA, Ueno T, Brumme ZL, Brockman MA. 2015. A robust and scalable TCR-based reporter cell assay to measure HIV-1 Nef-mediated T cell immune evasion. J Immunol Methods 426:104–13.

51. Trkola A, Matthews J, Gordon C, Ketas T, Moore JP. 1999. A cell line-based neutralization assay for primary human immunodeficiency virus type 1 isolates that use either the CCR5 or the CXCR4 coreceptor. J Virol 73:8966–74.

52. Lodge R, Lalonde JP, Lemay G, Cohen EA. 1997. The membrane-proximal intracytoplasmic tyrosine residue of HIV-1 envelope glycoprotein is critical for basolateral targeting of viral budding in MDCK cells. EMBO J 16:695–705.

53. Sauter D, Schindler M, Specht A, Landford WN, Munch J, Kim KA, Votteler J, Schubert U, Bibollet-Ruche F, Keele BF, Takehisa J, Ogando Y, Ochsenbauer C, Kappes JC, Ayouba A, Peeters M, Learn GH, Shaw G, Sharp PM, Bieniasz P, Hahn BH, Hatziioannou T, Kirchhoff F. 2009. Tetherin-driven adaptation of Vpu and Nef function and the evolution of pandemic and nonpandemic HIV-1 strains. Cell Host Microbe 6:409–21.

54. Klimkait T, Strebel K, Hoggan MD, Martin MA, Orenstein JM. 1990. The human immunodeficiency virus type 1-specific protein vpu is required for efficient virus maturation and release. J Virol 64:621–9.

55. Dave VP, Hajjar F, Dieng MM, Haddad E, Cohen EA. 2013. Efficient BST2 antagonism by Vpu is critical for early HIV-1 dissemination in humanized mice. Retrovirology 10:128.

56. Westervelt P, Gendelman HE, Ratner L. 1991. Identification of a determinant within the human immunodeficiency virus 1 surface envelope glycoprotein critical for productive infection of primary monocytes. Proc Natl Acad Sci U S A 88:3097–101.

57. Lodge R, Bellini N, Laporte M, Salahuddin S, Routy JP, Ancuta P, Costiniuk CT, Jenabian MA, Cohen EA. 2020. Interleukin-1beta Triggers p53-Mediated Downmodulation of CCR5 and HIV-1 Entry in Macrophages through MicroRNAs 103 and 107. mBio 11.

58. Keele BF, Giorgi EE, Salazar-Gonzalez JF, Decker JM, Pham KT, Salazar MG, Sun C, Grayson T, Wang S, Li H, Wei X, Jiang C, Kirchherr JL, Gao F, Anderson JA, Ping LH, Swanstrom R, Tomaras GD, Blattner WA, Goepfert PA, Kilby JM, Saag MS, Delwart EL, Busch MP, Cohen MS, Montefiori DC, Haynes BF, Gaschen B, Athreya GS, Lee HY, Wood N, Seoighe C, Perelson AS, Bhattacharya T, Korber BT, Hahn BH, Shaw GM. 2008. Identification and characterization of transmitted and early founder virus envelopes in primary HIV-1 infection. Proc Natl Acad Sci U S A 105:7552–7.

59. Lee HY, Giorgi EE, Keele BF, Gaschen B, Athreya GS, Salazar-Gonzalez JF, Pham KT, Goepfert PA, Kilby JM, Saag MS, Delwart EL, Busch MP, Hahn BH, Shaw GM, Korber BT, Bhattacharya T, Perelson AS. 2009. Modeling sequence evolution in acute HIV-1 infection. J Theor Biol 261:341–60.

60. Salazar-Gonzalez JF, Bailes E, Pham KT, Salazar MG, Guffey MB, Keele BF, Derdeyn CA, Farmer P, Hunter E, Allen S, Manigart O, Mulenga J, Anderson JA, Swanstrom R, Haynes BF, Athreya GS, Korber BT, Sharp PM, Shaw GM, Hahn BH. 2008. Deciphering human immunodeficiency virus type 1 transmission and early envelope diversification by single-genome amplification and sequencing. J Virol 82:3952–70.

61. Salazar-Gonzalez JF, Salazar MG, Keele BF, Learn GH, Giorgi EE, Li H, Decker JM, Wang S, Baalwa J, Kraus MH, Parrish NF, Shaw KS, Guffey MB, Bar KJ, Davis KL, Ochsenbauer-Jambor C, Kappes JC, Saag MS, Cohen MS, Mulenga J, Derdeyn CA, Allen S, Hunter E, Markowitz M, Hraber P, Perelson AS, Bhattacharya T, Haynes BF, Korber BT, Hahn BH, Shaw GM. 2009. Genetic identity, biological phenotype, and evolutionary pathways of transmitted/founder viruses in acute and early HIV-1 infection. J Exp Med 206:1273–89.

62. Kitchen SG, Bennett M, Galic Z, Kim J, Xu Q, Young A, Lieberman A, Joseph A, Goldstein H, Ng H, Yang O, Zack JA. 2009. Engineering antigen-specific T cells from genetically modified human hematopoietic stem cells in immunodeficient mice. PLoS One 4:e8208.

