## Supplemental Figures for "HIV-1 Vpu complements HLA-A downregulation by Nef and impairs TCR-mediated recognition of HIV-infected cells through distinct mechanisms"

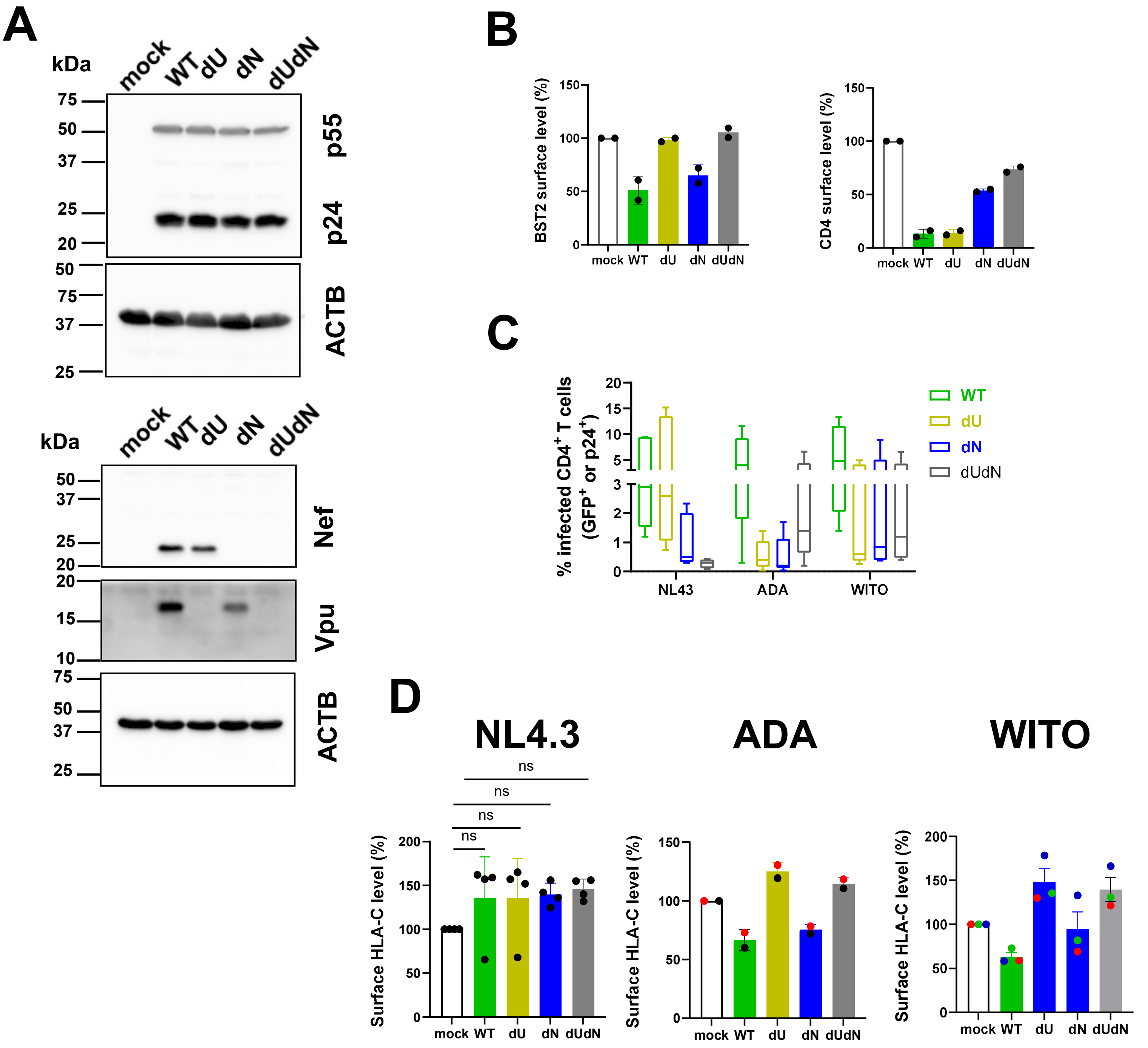


**Fig. S1: HIV-1 Vpu contributes to the modulation of class I HLA on infected CD4$^+$ T cells. Related to Fig. 1.** CD4$^+$ T cells from HLA-A*02:01 donors were infected for 72h with GFP-marked NL4.3, GFP-marked CCR5-tropic NL4.3-ADA (ADA), or WITO viruses expressing all viral proteins (wild-type, WT) or lacking Vpu (dU), Nef (dN) or both viral proteins (dU/dN). Uninfected cells (mock) were used as control. (**A-B**) Virus characterization in ADA-infected cells. Panel A: Western blots show expression profiles of proteins of interest following probing of infected cell lysates with the α-p24, α-Nef, α-Vpu and α-ACTB (actin) Abs, respectively. Panel B: Flow cytometry analysis shows expression levels of BST2 and CD4 (mean +/- SD) on infected GFP$^+$ cells. (**C-D**) Frequency of infected cells (n=5 donors) (Panel C) or level of HLA-C on infected cells (mean +/- SD) (Panel D) were determined using flow cytometry. In both Panels, infected cells were determined based on expression of GFP (for NL4.3 and ADA viruses) or p24 (for WITO virus). In Panels B, C and D, cells were stained with Abs against the indicated proteins, and surface levels for BST2, CD4 and HLA-C were determined by dividing the geomean fluorescence intensity (MFI) of the given marker on infected cells by that on bystander cells (GFP$^-$ or p24$^-$) and multiplied by 100. The value for mock-infected cells was given 100% based upon total cell gating. Each dot is one experiment, and the experiments with ADA, WITO viruses in Panel D were color-coded to highlight more clearly the effect of Vpu for a given donor. Statistical test: nonparametric Kruskal-Wallis with Dunn's multiple comparisons; ns, not significant.

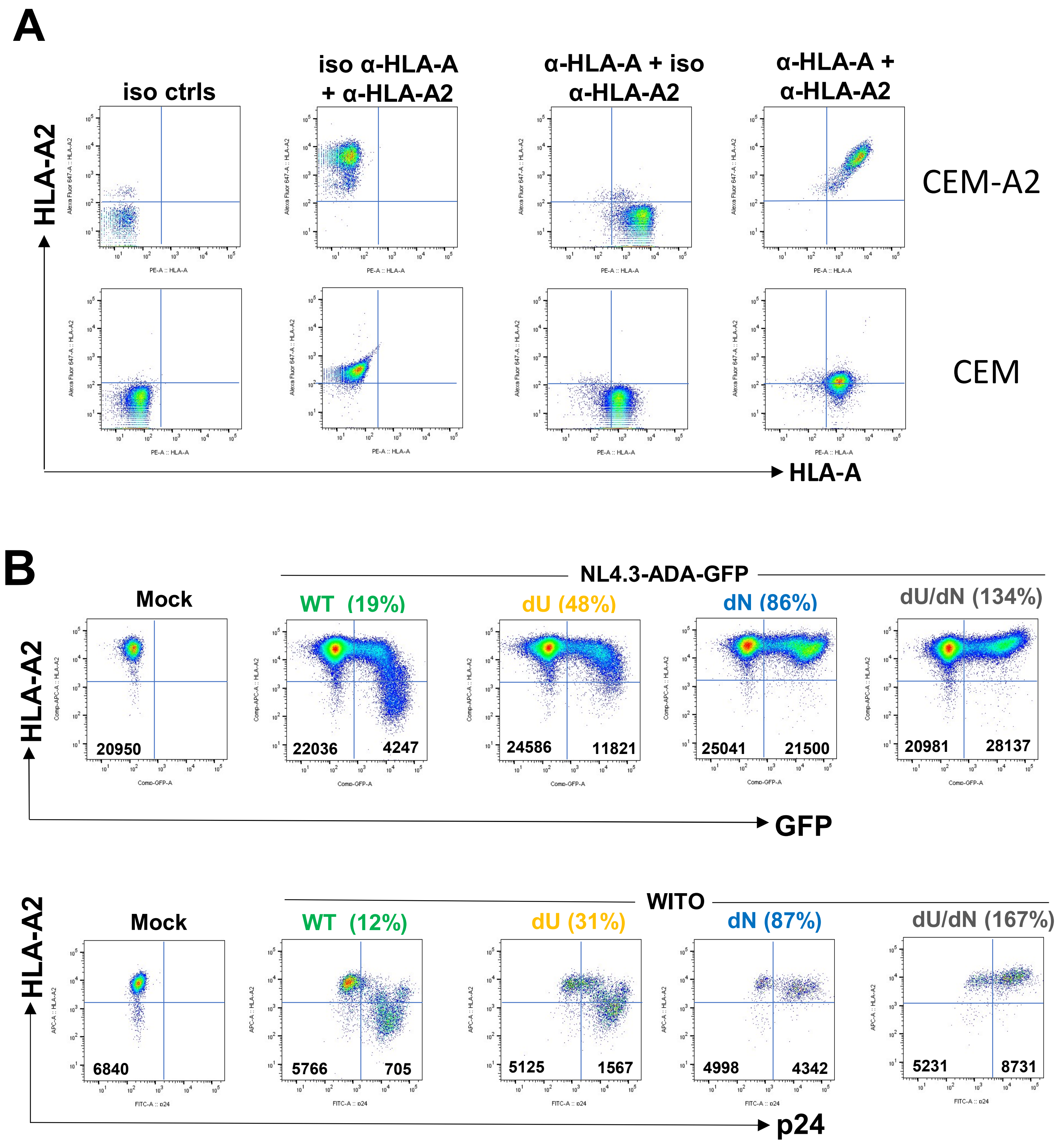


**Fig. S2: HIV-1 Nef and Vpu downregulate HLA-A2 expression on infected cells. Related to Figs 1 and 3.** (A) Parental CEM or HLA-A*02:01-expressing CEM (CEM-A2) were stained with Abs against HLA-A2, HLA-A or the corresponding isotype controls. The α-HLA-A Ab recognizes an epitope shared by HLA-A2, -A3, -A28, -A29, -A31 and -A33 alleles. Dot plots show that CEM-A2, but not CEM cells were strongly stained for HLA-A2. (B) CEM-A2 infected with NL4.3-ADA-GFP or WITO for 2 days and stained for surface HLA-A2 using the α-HLA-A2 Ab as described in Panel A. Flow dot plots showed % of HLA-A2 level (calculated as follows: geomean fluorescence intensity (gMFI) of HLA-A2 on infected cells was divided by that on uninfected cells x 100. Infected cells were defined as GFP$^{+}$ for NL4.3-ADA-GFP or p24$^{+}$ for WITO, while the uninfected was GFP$^{-}$ and p24$^{-}$, respectively. gMFI for infected and uninfected cells are as shown.

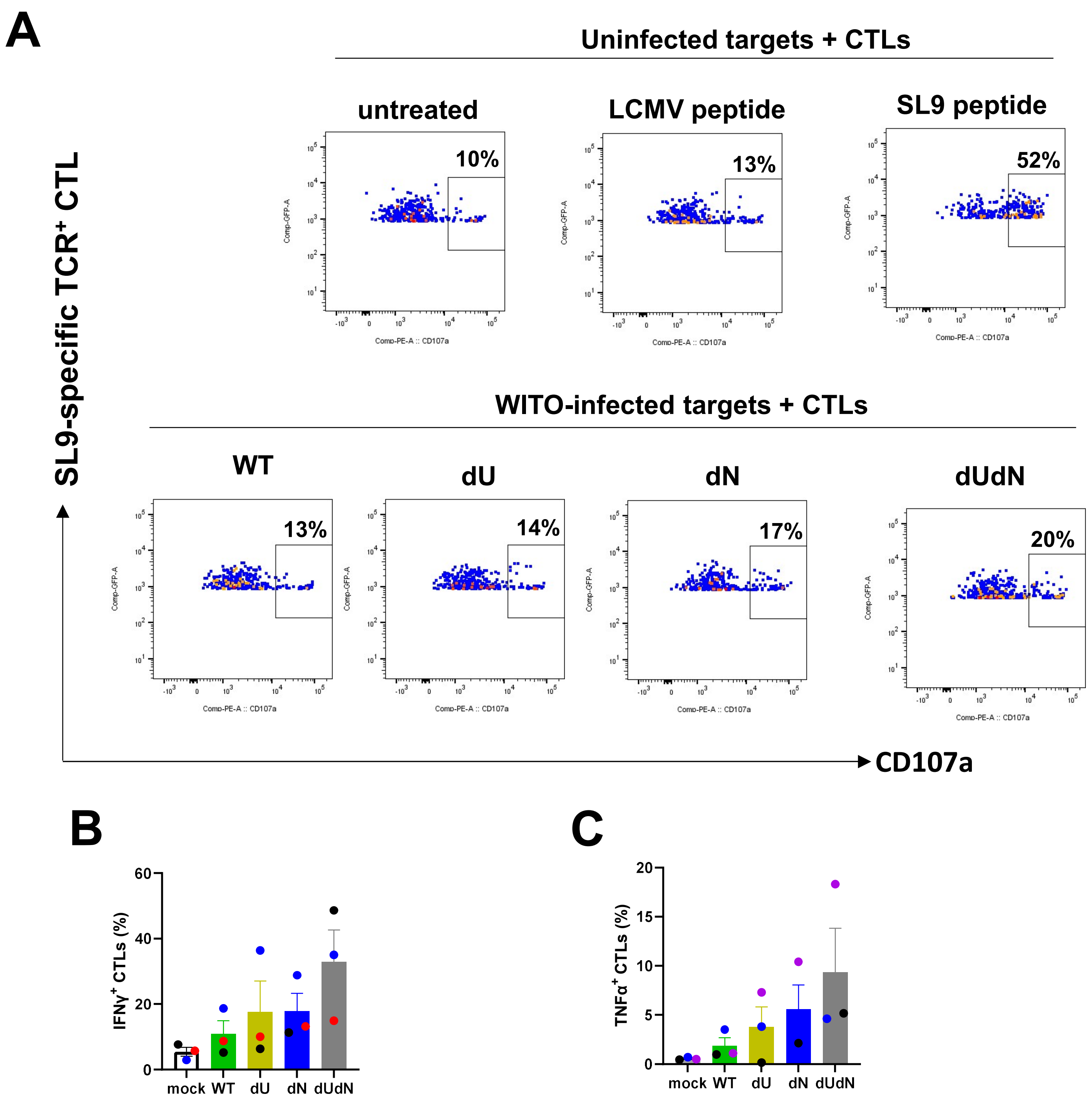


**Fig. S3: Downmodulation of cell-surface HLA-A2 is linked to potential reduction of HIV-specific CTL responses. Related to Fig. 3**. CEM-A2 CD4$^+$ T cells were infected with WITO viruses expressing all proteins (WT) or deficient of Vpu (dU), Nef (dN) or Vpu and Nef (dUdN) for 24 h. Alternatively, uninfected cells (mock) were untreated or pulsed with an LCMV peptide (GP276-286, 10 µg/ml) or HIV SL9 peptide (10 µg/ml) for 2h. The CEM-A2 cells (targets) were subsequently co-cultured for 3h with primary CD8$^+$ T cells (effectors) in the presence of PE-conjugated αCD107 Ab and monensin (added at hour 0). At the time of the co-culture, the effector cells had been transduced for 3-5 days with a GFP-marked HIV-based lentiviral vector expressing the HLA-A*02:01 restricted, SL9-specific TCR. The cytotoxic lymphocytes (CTLs) were analyzed by flow cytometry for the presence of CD107a at the cell surface (indicative of cell degranulation) and intracellular IFNγ and TNFα. For cytokine detection, Brefeldin A was added at hour 1 of the co-culture. At the end of the co-culture, cells were washed and successively stained with live/dead Zombie Aqua and α-CD8 Ab. Cells were fixed, permeabilized and stained for IFNγ and TNFα(A) An example of flow dot plots showing proportions of living (Aqua$^-$) GFP$^+$CD8$^+$ CTLs expressing CD107a. The data with uninfected targets pulsed with irrelevant (LCMV) or HIV-specific SL9 peptide confirms the HIV-specific CTL response. (B and C) Frequencies of Aqua$^-$ GFP$^+$CD8$^+$ IFNγ$^+$ CTLs and Aqua$^-$ GFP$^+$CD8$^+$ TNFα$^+$ CTLs over the different donors. Shown are mean +/- SD. Each color is a different donor.

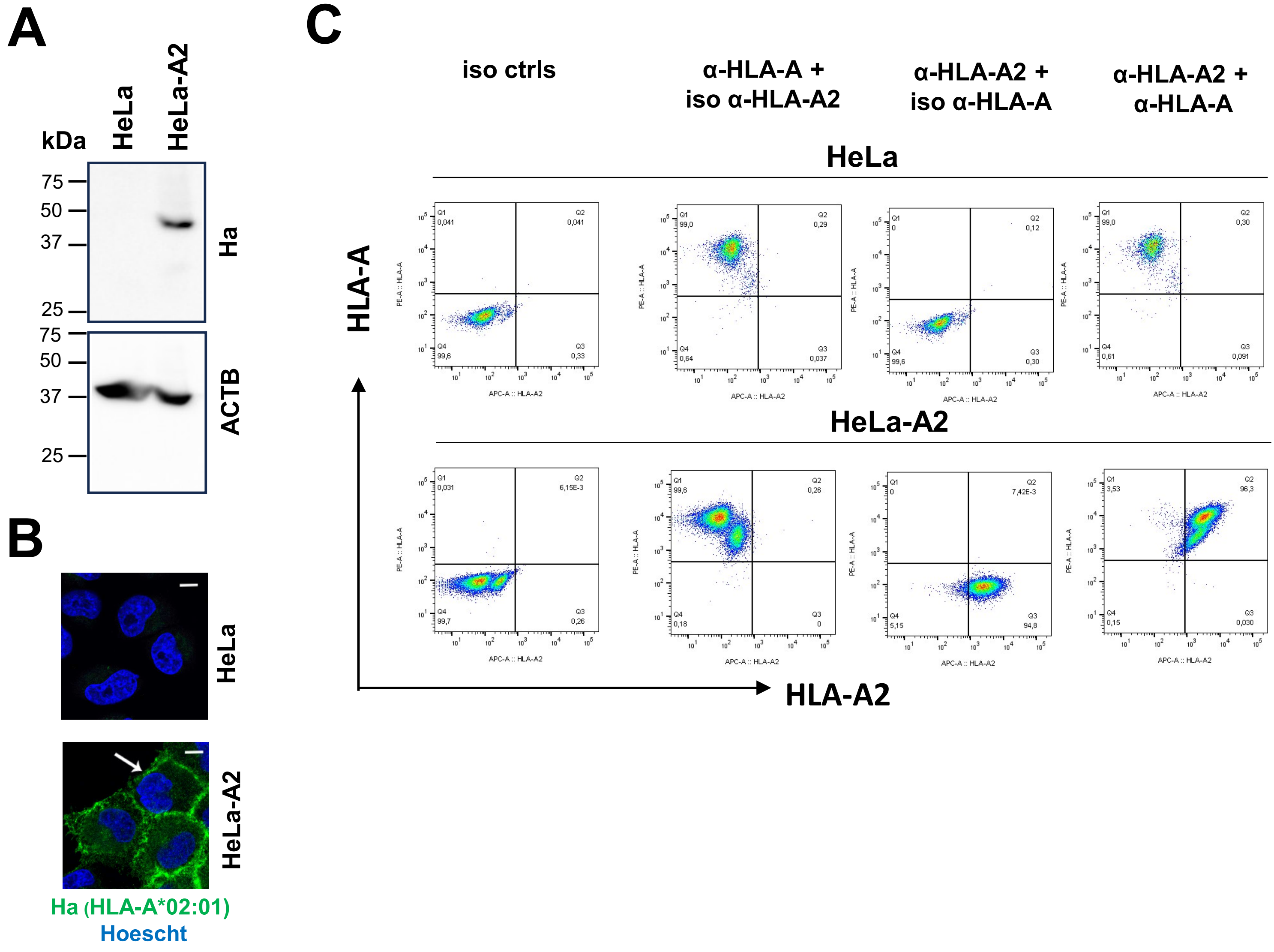


**FIG. S4. Characterization of HeLa-A2 cells that stably express Ha-tagged HLA-A*02:01. Related to Fig. 5**. (A) Western blot analysis with α-Ha Ab showing Ha-HLA-A2 detection in HeLa-A2 cells but not in the parental HeLa. (B) Confocal microscopy analysis reveals predominant staining of Ha-HLA-A2 at the surface of HeLa-A2 but not parental HeLa (white arrow). Hoechst counterstains cell nuclei, scale bar is 10 µm. (C) Flow cytometry analysis shows surface expression of HLA-A2 and HLA-A in HeLa-A2 cells compared to parental HeLa. Cells were co-stained with α-HLA-A2 and α-HLA-A Abs or isotype controls as described for Fig. S2A. The data reveal that parental HeLa cells were positive for HLA-A but negative for HLA-A2 while the HeLa-A2 cells were positive for both proteins with the HLA-A2 seemingly stained less strongly than the endogenous non-HLA-A2 alleles.

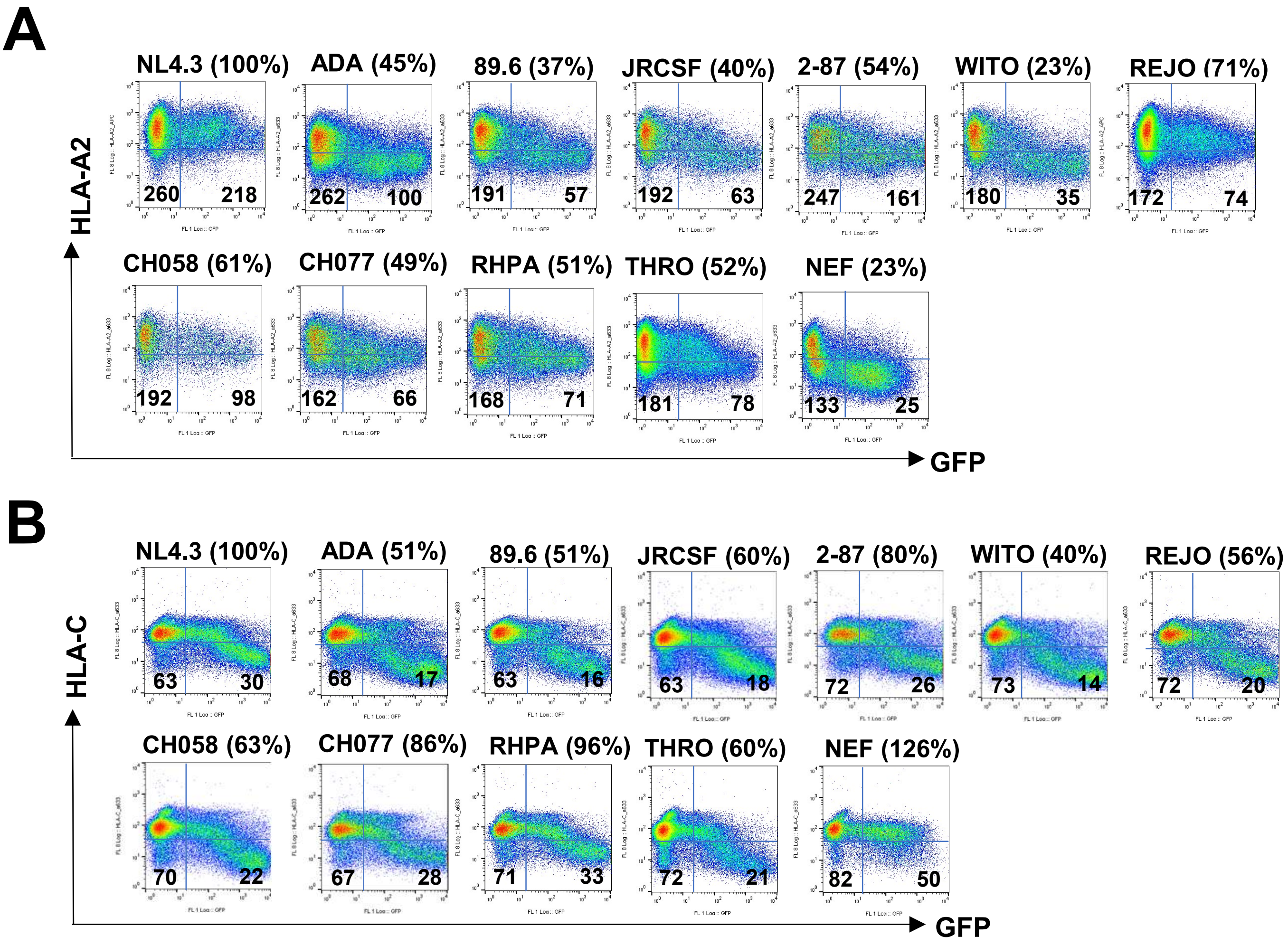


**FIG. S5. HIV-1 Vpu alone is sufficient to modulate HLA-A2 expression. Related to Fig. 5.** (A-B) Representative flow graphs showing HLA-A2 and HLA-C expression on HeLa-A2 cells that were transfected for 48 h with a GFP-marked plasmid expressing Vpu or Nef (control) and stained with α-HLA-A2 and α-HLA-C Abs, respectively. In all Panels, expression levels in percentages were determined as follows. gMFI of the indicated protein on transfected ($GFP^+$) cells was divided by that on the corresponding untransfected $GFP^-$ cells, and the ratio obtained with the NL4.3 Vpu was set at 100% (reference). HLA-A2 or HLA-C levels on cells transfected with the other Vpu/Nef plasmids were expressed relative to the reference.

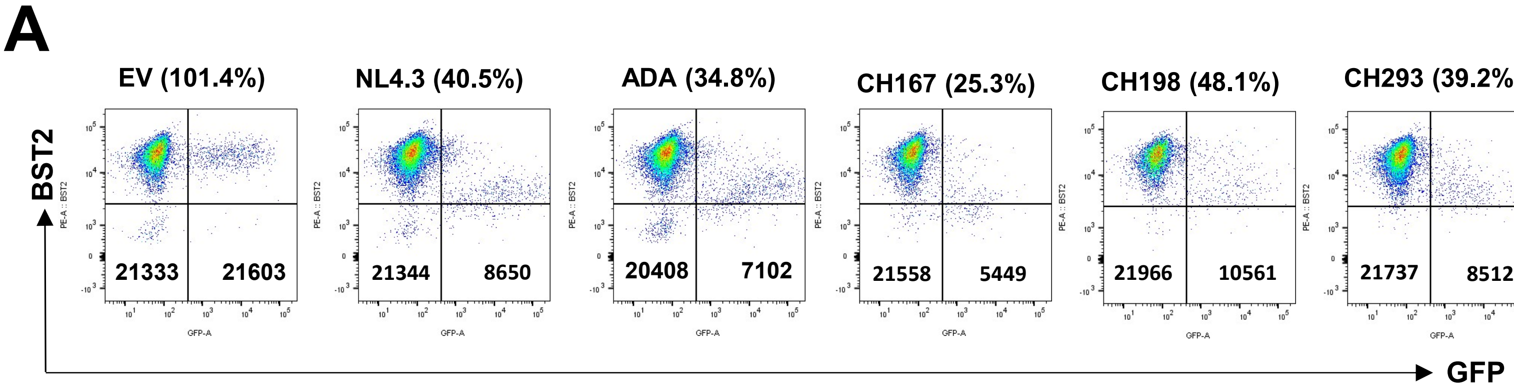


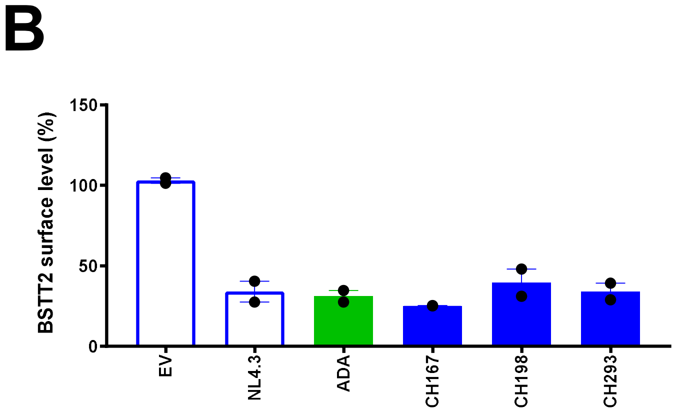


**FIG. S6. The inability of CH-293 Vpu to downregulate HLA-A2 expression is not due to expression defect. Related to Fig. 5**. Hela-A2 were transfected for 48 h with a GFP-marked plasmid expressing no Vpu (empty vector, EV) or Vpu. Transfected cells were examined for BST2 expression by flow cytometry using an α-BST2 Ab. Expression levels in percentages were determined as follows. gMFI of BST2 on transfected ($GFP^+$) cells was divided by that on the corresponding untransfected $GFP^-$ cells. (A) Representative flow graphs showing BST2 expression (gMFI) on $GFP^+$ and $GFP^-$ cells. (B) BST2 expression (mean +-/ SD) on transfected cells in two independent experiments.

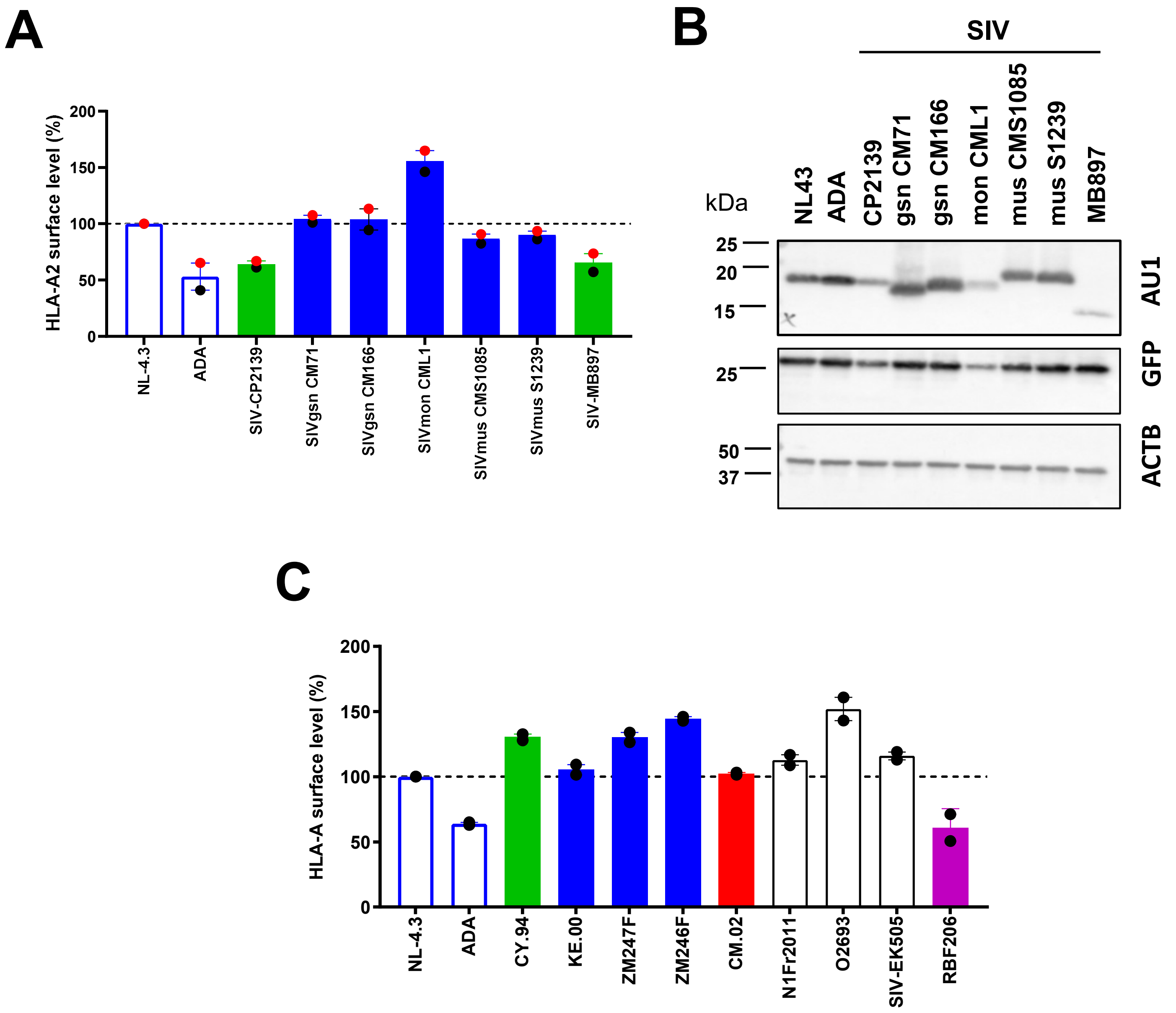


**FIG. S7. Vpu gains HLA-A modulating function likely precedes emergence of SIVcpz and pandemic HIV-1. Related to Fig. 5**. HeLa-A2 (Panels A and B) or parental HeLa (Panel C) were transfected with a GFP-harbouring plasmid expressing AU1-tagged Vpu from different primary viruses. At 48 h post transfection, cells were analyzed by flow cytometry for surface HLA-A2 or HLA-A using the α-HLA-A2 or α-HLA-A Ab, respectively. Shown is cell surface expression on transfected ($GFP^+$) cells relative to that on NL4.3 Vpu-expressing cells. The latter was set at 100%. Panel B: Western blotting analysis of cells lysates used in the experiments shown in Panel A. Protein lysates were probed for AU1 (Vpu), GFP (transfection efficiency) or actin (ACTB, loading control). Each dot is an individual experiment and the results were expressed as mean +/- SD.

| | | | -A2 | -C |
|---|---|---|---|---|
| MB_ADA | M-----QPLQILAIVALVVAAIIAIVVWTIVF | 27 | ++ | ++ |
| MB_89.6 | M-----LSLQILAIVALVVAAIIAIVVWSIVF | 27 | ++ | ++ |
| MB_WITO | M-----QPLEILAVVALVVALILAIVVWTIVY | 27 | ++ | ++ |
| MC_CH167 | MFDFIARVDYRVGVVALVVALIIAIIVWSIVY | 32 | ++ | ++ |
| MC_CH198 | MLAFLARVDYRLGVGALVIAGIIAIVVWIIAY | 32 | ++ | ++ |
| MD_UG.99 | M-----QPLEIGAIVALVVALIIAIVVWTIVF | 27 | ++ | ++ |
| MD_ZA.90 | M-----QPLVILAIVALVVALILAIVVWTIVF | 27 | ++ | ++ |
| O_RBF206 | M-----HQRDVLVLVVFIALSIVSLLIWLFSL | 27 | ++ | ++ |
| MB_JRCSF | M-----QPLQILAIVALVVAGIIAIIVWSIVL | 27 | ++ | + |
| MA_SE.95 | M-----TPLEIWAIVGLVVVSIIAIVVWTIVG | 27 | ++ | +/- |
| MB_REJO | M-----QTLQILAIVALVIAGIIAIVVWTIVF | 27 | + | ++ |
| [M]SIV_MB897 | M-----EIFIILGLIGIVIELVIAIVVWLKAY | 27 | + | + |
| [O]SIV_2139 | M-----HPRDIIVIIIGITLLAVTVIIWLKIF | 27 | + | + |
| MB_NL4.3 | M-----QPI-IVAIVALVVAIIIAIVVWSIVI | 26 | - | - |
| MB_2_87 | M-----KSLETLAIVALVVAAILAIVVWSIVF | 27 | +/- | + |
| MB_CH058 | M-----QPLN-IAIGALIVAAILAIIVWTIVF | 26 | +/- | + |
| MB_THRO | M-----SPLVIASIVALVVAAIIVIVIWSIVF | 27 | +/- | + |
| MF_CM.02 | M-----SYLIIILVIVAFIVALIAAIIVWTIVY | 27 | +/- | +/- |
| MB_CH077 | M-----QSLYILGIVALVVAAILAIVVWTIVY | 27 | + | +/- |
| [N]SIV_EK505 | MLLL-----IKLGFIGLAIETLIVIVVWAIVY | 27 | +/- | +/- |
| O_HJ100 | M-----QHKE--LLIIICALLLINVMLWMFIL | 25 | - | +/- |
| O_HJ162 | M-----HQKDLLLLIVIIALLLTNVILWLFNL | 27 | +/- | +/- |
| MB_RHPA | M-----QSLQIVAIVAVVVGIIIAIVVWSIVL | 27 | +/- | - |
| MA_CY.94 | M-----LPLVILAIVGLIVALILAIVVWTIVF | 27 | - | - |
| MC_KE.00 | MLDLLARVDYRLGVGALIVALIIAIIVWTIVY | 32 | +/- | +/- |
| MC_ZM247 | MLEL----DYKIAIAALIVALIIAIVVWTIVY | 28 | - | - |
| MC_ZM246 | MISLIEKVDYRLGVGALIVALIIAIVVWTIAY | 32 | - | - |
| MC_CH293 | MLSWIEKADYRLGVGALVVAIIIAIIVWIIAY | 32 | - | - |
| N_N1FR2011 | MLLL--------GFIAVGVAFIIAVVIWVLLY | 24 | - | - |
| N_CK1.62 | MLLL--------GFIAVGIAIVIAVIIWVLLY | 24 | - | - |
| N_2693 | MLLL--------GFIAVGIAIVIAAIIWVLLY | 24 | - | - |
| P_RBF168 | M-----HPRDEAVLIIAGVLLICIIVVWGKVL | 27 | - | - |
| P_06OCMU | M-----HPRDEAVLIIAGVLLLCCIVXWGKVL | 27 | +/- | - |

MB- group M, clade B

**FIG. S8. Sequence alignment of the N-terminal/transmembrane region (TM) of Vpu variants studied. Related to Figs. 5 and 6**. The alignment was generated using the Clustal Omega program. Vpu ability to downregulate HLA-A2 or HLA-C and the extent of the downregulation are denoted by plus (+) and minus (-) signs based on the transient expression experiments with HeLa cells such as those shown in Fig. 5. The ++ sign indicates 40-60% reduction compared to the control NL4.3 Vpu. The + sign indicates less than 40% reduction compared to the control NL4.3 Vpu. The +/- denotes variable detection of activity in different experiments. The length of the region is indicated at the end of the alignment. Amino acid identities that are not frequently found or absent when compared to the active Vpu alleles (the first group) are highlighted in pink.

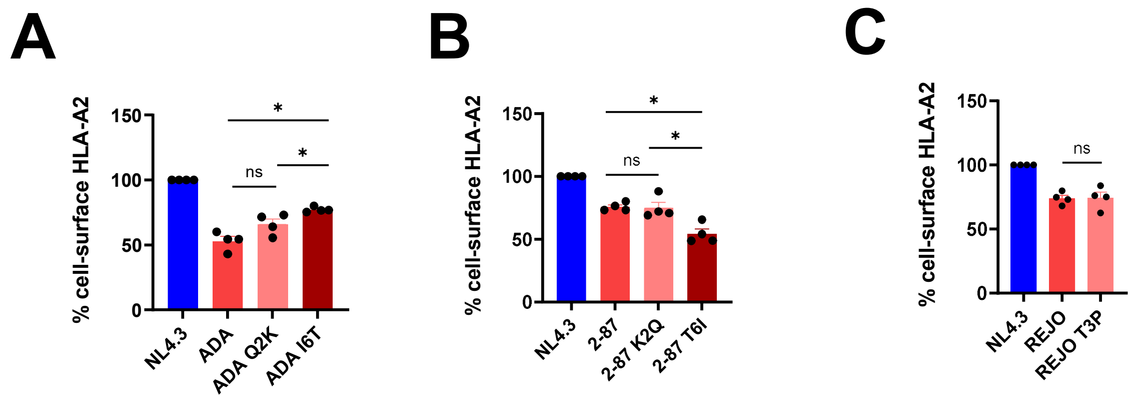


**FIG. S9**. **Assessing the importance of individual residues in the N-terminus/TMD in Vpu ability to downregulate HLA-A2**. **Related to Fig. 6.** HeLa-A2 cells were transfected with a GFP-harbouring plasmid expressing Vpu in which different single point mutations were made in Vpu from (**A**) ADA, (**B**) 2-87 or (**C**) REJO. In Panel A, the amino acids at indicated positions in ADA Vpu were swapped for those in 2-87 Vpu, and conversely for Panel B. In Panel C, the amino acids at position 3 in REJO Vpu was swapped for that in ADA Vpu or NL4.3 Vpu. At 48 h post transfection, cells were analyzed by flow cytometry for surface HLA-A2 using the α-HLA-A2 Ab described in the paper. HLA-A2 expression on transfected ($GFP^+$) cells is as indicated, and expression levels were determined as described for Fig. S2. Statistical test: Mann-Whitney's U-test; ns, not significant. * $p$=0.028.

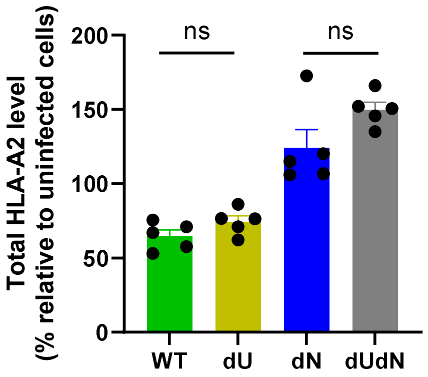


**Fig S10: Vpu does not affect the total level of HLA-A2 in infected CD4+T cells. Related to Fig. 9**. Primary CD4$^{+}$T were infected with NL4.3-ADA-GFP (WT or mutant viruses lacking Nef and/or Vpu) for 2 days, then fixed, permeabilized and analyzed for total (surface and intracellular) HLA-A2 by flow cytometry using the α-HLA A2 Ab. HLA-A2 levels (mean +/- SD) were determined as follows: gMFI of HLA-A2 of infected GFP$^{+}$ cells was divided by that of uninfected GFP$^{-}$ cells x 100. Each dot is a donor. Statistical test: nonparametric Mann-Whitney's; ns, not significant.
